# Oncostatin M cytokine promotes breast cancer progression by remodelling the extracellular matrix and activating integrin signalling in cancer cells

**DOI:** 10.64898/2026.06.01.729048

**Authors:** Peio Azcoaga, Andrea Abaurrea, Uxue Alvarez-Huesa, Paula Duch, Angela M. Araujo, Joanna I. López-Velazco, Zuhara Telletxea, Marta Rezola, Juana M. Flores, Gerhard Müller-Newen, Ana M Aransay, Mikel Azkargorta, Felix Elortza, David Otaegui, Steve Stegen, Jai Prakash, Sara Manzano, María M. Caffarel

## Abstract

Tumours reshape their surrounding extracellular matrix (ECM), creating a microenvironment with altered chemical and mechanical properties. Integrins detect these changes, linking the ECM to the intracellular cytoskeleton and promoting cell survival, motility, invasion and differentiation, and further ECM remodelling. However, the molecular mechanisms by which tumours remodel their ECM are not well understood. Here, we found that the cytokine oncostatin M (OSM) promotes breast cancer progression by activating ECM remodelling and integrin signalling in cancer cells, as shown by combining complementary *in vitro*, *in ovo* and *in vivo* models, and transcriptomic and proteomic analyses. We demonstrated that OSM induces fibrosis, characterized by increased collagen deposition and hydroxylation, together with activation of ECM and ECM-associated proteins and modifiers such as fibronectin, tenascin C, LOX, PLOD2 and collagen prolyl hydroxylases. OSM also promoted the expression of integrins. Integrin alpha 5 (ITGA5) was identified as an important mediator of OSM-effects. ITGA5 blockade, by means of small interference RNA and therapeutic inhibition with a blocking peptide, abrogated OSM-induced cancer cell migration, invasion and *in vivo* tumour growth. In addition, OSM blockade with a specific inhibitor reduced tumour growth in an immunocompetent mouse model. Our results are clinically relevant as the expression of integrins and matrisome genes strongly correlated with *OSM* and its receptor *OSMR* in breast cancer clinical samples; and co-expression of *OSMR* and *ITGA5* associated with decreased survival in basal breast cancer patients. Collectively, our data reinforce the potential of the OSM-ITGA5 axis as a therapeutic target in this breast cancer subtype, which shows the highest mortality rates.

## Introduction

Tumours actively modify and accumulate extracellular matrix (ECM) components, generating a tumour-supportive microenvironment with unique chemical and mechanical properties^1^. Integrins, the main cell-surface adhesion receptors, detect these ECM alterations by coupling matrix molecules to the actin cytoskeleton^2^. These interactions transmit bidirectional forces that activate signalling pathways involved in cell survival, differentiation, migration, and continued ECM remodelling. The activation of this dynamic machinery is fundamental to disrupt physical barriers and promote cancer cell dissemination during early metastasis, as well as for the colonization and outgrowth in the target organ^3^. Fibroblasts are the main ECM producers, and besides their functions in ECM deposition and remodelling, they can favour angiogenesis and regulate immune responses by secreting a plethora of cytokines^4^. Importantly, ECM can also be produced by cancer cells^1,5^ but the mechanisms by which tumour cells remodel their ECM are not fully characterized.

Here we show that the cytokine oncostatin M (OSM) promotes breast cancer progression by remodelling the ECM and activating integrin signalling. Cytokines are often dysregulated in cancer, and have been linked to all stages of cancer, from tumour initiation to metastasis, as they mediate the crosstalk between different cell types within the tumour microenvironment^6,7^. OSM is an understudied pleiotropic cytokine belonging to the interleukin-6 (IL6) family, which is considered one of the most important cytokine families in linking inflammation and cancer^8,9^. OSM is mainly secreted by myeloid cells and binds to its main specific receptor OSMR, which is expressed by a wide range of cell types including mesenchymal, mesothelial, endothelial and epithelial cells. OSM has multiple functions in haematopoiesis, liver regeneration, wound healing and metabolism, among others^10,11^. Its dysregulation has been linked to inflammatory diseases, fibrosis and cancer. We and others have demonstrated that OSM promotes cancer progression in preclinical models of glioblastoma, breast, gastric, pancreatic and cervical cancer^12–16^. However, the molecular mechanisms by which OSM exerts its pro tumour effects have been poorly investigated.

Our data demonstrate that OSM promotes ECM deposition, collagen hydroxylation and integrin expression in cancer cells *in vitro* and *in vivo,* with or without the presence of cancer-associated fibroblasts (CAFs). It also shows that integrin alpha 5 (ITGA5) is an important mediator of OSM’s pro-malignant effects. We analysed the expression of OSM-OSMR-ITGA5 pathway in different breast cancer subtypes and identified basal (or triple-negative breast cancer, TNBC) as the most suitable patient subset to benefit from anti-OSM therapies and/or ITGA5 inhibitors. OSM and ITGA5 inhibition with therapeutic peptides reduced tumour growth in in vivo preclinical models of TNBC. In summary, our results support the potential of the OSMR-ITGA5 axis as a therapeutically actionable target in this breast cancer subtype.

## Materials and methods

### Reagents

References for all reagents used are described in **Supplemental Table 1**.

### Cell culture

Human breast cancer–associated fibroblasts 173 (CAF-173)^17^ were cultured in collagen-precoated flasks (Corning). Murine (4T1) and human (MDA-MB-231 and SK-BR-3) breast cancer cell lines were purchased from American Type Culture Collection (ATCC) and cultured following ATCC instructions. All cell lines were authenticated by short-tandem-repeat profiling (Genomics Core Facility at “Alberto Sols” Biomedical Research Institute) and routinely tested for mycoplasma contamination. Recombinant human OSM (rhOSM) or murine OSM (rmOSM) (R&D Systems) were added to cells at 10 ng/ml unless otherwise specified. For experiments combining recombinant murine OSM and OSM inhibitor (iOSM)^18,19^ in 4T1 cells, 5 ng/ml rmOSM and multiple concentrations of iOSM were used ranging from 1 to 40 iOSM:OSM molar ratios, equivalent to 30 to 1200 ng/ml of iOSM. 3D heterospheroids composed of MDA-MB-231 breast cancer cells and CAFs cells were grown in non-treated U-bottom 96-well plates (Corning), coated with 100 μl of 1% (w/v) Pluronic water (Pluronic F127, Thermo Scientific) 24 h prior to cell seeding to avoid cell-plate adhesion. Plates were rinsed with ultra-pure water (Invitrogen) before cell seeding.

Breast cancer cells and CAFs heterospheroids were seeded with 2.5 % basement membrane matrix (Geltrex A14133-02) in a 1:5 ratio (1000 breast cancer cells, 5000 CAFs and 1.25 μl matrix in 50 μl of Dulbecco’s Modified Eagle’s Medium (DMEM) per well). 72h later, they were treated with 30 ng/ml rhOSM or vehicle and treatment was repeated every 2-3 days. Pools of spheroids (6-12 per condition) were collected on day 10 after two centrifugations at 300 x g for 5 min and snap-frozen in liquid nitrogen.

### Cell transfections

Control and OSM-overexpressing MDA-MB-231 cells were generated as previously described^13^. OSMR-overexpressing and control SK-BR-3 cells were generated by transfection with pcDNA3.1/Zeo-OSMR or the empty vector pcDNA3.1/Zeo plasmids respectively and selected with 300 μg/ml of zeocin. Plasmids were kindly provided by Prof. Nicholas Coleman (Cambridge University, UK). Human ITGA5 was silenced with small interference RNAs (siRNAs), in parallel to its corresponding negative control siRNA non-targeting pool (siNTP, Dharmacon; **Supplemental Table 2**). MicroRNA (miRNA) transfections were performed using a synthetic hsa-miR-148a-3p miRNA and a negative control (scramble, scr) from the miRIDIAN miRNA Mimic Library (Horizon Discovery; Supplemental **Table 2**). All transfections were performed with Lipofectamine RNAiMax (Invitrogen) following manufacturer’s instructions. Cells were used for experiments 24h after transfection.

### Cell migration and invasion assays

MDA-MB-231 cells were subjected to migration and invasion assays for 48 hours by seeding 25.000 cells at the top of clear (for migration assays) or matrigel-coated (for invasion assays) 24-well transwell inserts respectively (8 μm pore, Corning). Foetal bovine serum (FBS) was used as a chemoattractant. Chambers were fixed in 10% formalin (20 minutes) and stained with crystal violet solution (20 minutes). For the quantification of migrated cells, crystal violet was solubilized with 600 μL of 1% SDS (30 minutes) and absorbance was measured at 570 nm.

### Mouse studies

In orthotopic mouse experiments, cell suspensions were injected in 50 µl serum-free and antibiotic-free DMEM. Cells were injected into the fourth right mammary fat pad of anaesthetized (with 4% isoflurane) 6- to 8-week-old female mice purchased from Charles River. For the syngeneic 4T1 orthotopic tumour experiment, 5.000 viable murine parental 4T1 cells were injected into syngeneic BALB/C mice (Charles River) and therapeutic neutralization of OSM was performed with a murine OSM inhibitor (iOSM), consisting on an Fc-tagged soluble OSMR–gp130 fusion protein^18,19^. For iOSM treatment, animals harbouring 4T1 tumours were treated intraperitoneally (i.p.) with iOSM or the control peptide (150 µg/animal) 3 times a week for 2 weeks, starting when tumours became detectable by palpation. For the MDA-MB-231 xenograft experiments, 100.000 viable control or OSM overexpressing (OSM) human MDA-MB-231 breast cancer cells, with or without 500.000 viable CAF-173 cells, were orthotopically injected into the mammary gland of athymic Nude-Foxn1 mice (Envigo). In the case of co-injections, cells were injected in 50% of DMEM medium and 50% of growth factor reduced (GFR) matrigel (Corning). When indicated, animals were treated (i.p.) with 20 mg/kg AV3 (ITGA5 inhibitor^20^) or vehicle (PBS, Corning) with DMSO (Sigma) 3 times a week starting when tumours reached around 80 mm^3^ in size. Animals were monitored by palpation 3 times a week and tumour growth was measured using a calliper. Tumour volume was calculated as (4π/3) × (width/2)2 × (length/2). Animals were culled once tumours reached the maximum allowed size.

After animal culling, lungs were visually inspected for macroscopic metastases. Tumours and lungs were divided into portions for (a) H&E (Sigma-Aldrich) and picrosirius red (Polysciences) pathological analyses (fixed in neutral buffered formalin solution (Sigma-Aldrich)) and (b) molecular analyses (snap frozen).

### Inoculation of tumour cells in the chorioallantoic membrane of fertilized chicken eggs (*in ovo* CAM model)

This model of tumorigenesis was performed as previously described^21^. 2x10^6^ MDA-MB-231 breast cancer cells (OSM-overexpressing vs. control, or siITGA5 OSM-overexpressing vs. siNTP OSM-overexpressing) were diluted in GFR matrigel (1:1 ratio in DMEM medium, Corning) and injected in each egg. After 7 days, tumours were excised, weighted and snap frozen.

### *Alu* quantification in genomic DNA

Genomic DNA (gDNA) was extracted from frozen lungs or distal CAM using the QIAmp DNA mini kit (Qiagen) for qPCR analysis of human *Alu* sequences normalising either with murine or chicken housekeeping gene expression^22^ (**Supplemental Table 2**).

### Western blotting

Samples were lysed in RIPA buffer (Millipore) supplemented with protease and phosphatase inhibitors (complete ULTRA Tablets, Mini, EASYpack protease inhibitor cocktail; and PhosSTOP phosphatase cocktail, both from Roche). Total lysates were quantified by bicinchoninic acid assay (BCA) (Pierce BCA Protein Assay Kit, Thermo Fisher Scientific), resolved by SDS-PAGE, and transferred to nitrocellulose membranes. After blocking with 5% (w/vol) non-fat dry milk (Millipore) in TBS-Tween, membranes were incubated with the corresponding antibodies (**Supplemental Table 3**) overnight at 4°C. Secondary antibodies (**Supplemental Table 3**) were chosen according to the species of origin of the primary antibodies and detected using ECL prime detection reagent (Invitrogen) in an iBright CL750 Imager (Invitrogen). Densitometric analysis of the bands was performed with Fiji ImageJ software.

### Measurement of hydroxyproline content

Collagen proline hydroxylation was quantified using a colorimetric protocol as previously described ^23^. Cultured cells or tumour tissue were hydrolysed for 3.5 hours at 135°C in 6 N HCl. Samples were then vacuum-dried and resuspended in deionized water. Hydroxyproline residues were oxidized with 0.05 M chloramine-T (Sigma-Aldrich), followed by the addition of Ehrlich’s aldehyde reagent (10% w/v p-dimethylaminobenzaldehyde dissolved in a mixture of n-propanol and perchloric acid (75:25, v/v); all from Sigma-Aldrich) and incubated for 45 minutes at 65°C to allow chromophore development. The absolute hydroxyproline content was determined using a standard curve generated with hydroxyproline standard (Sigma-Aldrich) and normalized to tissue weight or to the protein content of a parallel sample as measured by BCA assay.

### RNA extraction and RT-qPCR

RNA was obtained from snap frozen animal tissue or cell pellets and extracted using TRIzol reagent (Invitrogen) for RT-qPCR, microarray and RNA-seq gene expression analysis. cDNA was obtained with the Maxima First-Strand cDNA synthesis kit (Thermo Fisher Scientific) with DNAse treatment incorporated. RT-qPCR was performed using Power SYBR Green PCR master mix (Applied Biosystems).

Primer sequences are indicated in **Supplemental Table 2** and were purchased from Condalab. miRNA expression was detected using the Taqman technology in which miRNAs from 100-500 ng of total RNA were converted to cDNA by using the TaqMan MicroRNA Reverse Transcription Kit (ThermoFisher Scientific). The following PCR program was used: 30 min at 16°C, 30 min at 42°C and 5 min at 85°C. RT-qPCR was performed combining 2-4 ng of cDNA with the TaqMan MicroRNA Assays (**Supplemental Table 2**) and the TaqMan Universal Master Mix II, no UNG (Thermo Fisher Scientific). The RT-qPCR was performed under the following conditions: 10 min at 95°C followed by 40 cycles of 95°C for 15 sec and 60°C for 1 min. Expression levels of genes and miRNAs were determined using the ΔΔCt method^24^ and normalized against housekeeping sequences optimized for each reaction^25^.

### Transcriptomic analyses

Microarray analysis of control and OSM-activated MDA-MB-231 cells and derived orthotopic tumours was performed using the Human Clariom S assay (Thermo Fisher Scientific). RNA quality was evaluated using the 2100 Bioanalyzer (Agilent) and microarray chips were processed on the Affymetrix GeneChip Fluidics Station 450 and Scanner 3000 7G (Affymetrix) according to standard protocols (n = 3 per experimental condition). Data were analysed using the Transcriptome Analysis Console 4.0 (TAC). Genes with false discovery rates (FDR)<0.05 (*in vitro*) or 0.1 (*in vivo*) and absolute fold change (FC>2) were considered significantly modulated. RNA-seq analysis was performed in frozen samples of control and OSM-activated orthotopic tumours derived from co-injection of MDA-MB-231 and CAF-173 cells. For RNA-seq library preparation, RNA sample quantity and quality were measured using Qubit RNA HS Assay Kit (Thermo Fisher Scientific) and Agilent RNA 6000 Nano Chips (Agilent Technologies), respectively. cDNA was synthesized with SuperScript-II Reverse Transcriptase (Thermo Fisher Scientific), and libraries were prepared using TruSeq Stranded mRNA library Prep kit (Illumina Inc.) and TruSeq RNA CD Index Plate (96 Indexes, 96 samples; Illumina Inc.), following TruSeq Stranded mRNA Sample Preparation Guide (part no. 15031058 Rev. E). The final dsDNA libraries were quantified by Qubit dsDNA HS DNA Kit (Thermo Fisher Scientific) and qualified in an Agilent 2100 Bioanalyzer using Agilent High Sensitivity DNA kit (Agilent Technologies). Paired-end Illumina sequencing was performed. For the RNA-seq analysis, sequencing data were converted into raw data (FASTQ files) using the Illumina bcl2fastq Conversion Software. The raw sequence reads were aligned to the Human Reference build number 38 (hg38) using STAR^26^. Read counts were obtained for each gene using featureCounts v2.0.3^27^. Differential gene expression analysis was performed by the *DESeq2* R package^28^. Genes with FDR<0.05 and absolute FC>2 were considered significantly modulated.

### Proteomics

Proteomic analyses were performed in frozen samples of control and OSM-activated orthotopic tumours derived from co-injection of MDA-MB-231 and CAF-173 cells. Samples were processed by the Proteomic Platform Service at CIC bioGUNE (Derio, Spain) for the relative quantification of the proteins by label free LC-MS/MS, following the protocol described by Wisniewski et al^29^ with minor modifications, and analysed in a hybrid trapped ion mobility spectrometry – quadrupole time of flight mass spectrometer (timsTOF Pro with PASEF, Bruker Daltonics) coupled online to a nanoElute liquid chromatograph (Bruker). Protein identification and quantification were carried out using PEAKS X software (Bioinformatics solutions)^30^. Searches were carried out against a database consisting of *Homo sapiens* and *Mus musculus* entries from Uniprot Swissprot^31^. Oxidation of methionines and proline hydroxylation were considered as variable modifications, whereas carbamidomethylation of cysteines was considered as fixed modification.

Precursor and fragment tolerances of 20 ppm and 0.05 Da were considered for the searches, respectively. Only proteins identified with at least one peptide at FDR<1% were considered for further analysis. Protein abundance was compared by means of a student’s *t*-test, being proteins with a *P* value<0.05 considered as significantly deregulated.

### Gene pathway analyses

The functional classification of the statistically significant differentially expressed genes and proteins was performed using the ENRICHR tool (https://maayanlab.cloud/Enrichr/)^32^. REVIGO tool^33^ (http://revigo.irb.hr/) was used to reduce gene ontology (GO) analyses by removing redundant GO terms. Gene set enrichment analysis (GSEA) was performed as previously described^34^ using GSEA software from the BROAD Institute. Gene signatures analysed are indicated in **Supplemental Table 4**. For GO pathway and GSEA analyses, FDR<0.05 were considered statistically significant. Ingenuity® pathway analysis (IPA, QIAGEN) was also used for a characterization of biological processes in murine proteomic and transcriptomic data.

The calculated *P* values for the different analyses performed determine the probability that the association between proteins in the dataset and a given process, pathway or upstream regulator is explained by chance alone, based on a Fisher’s exact test (*P* value <0.05 being considered significant). The activation z-score represents the bias in gene regulation that predicts whether the upstream regulator exists in an activated (positive values) or inactivated (negative values) state, based on the knowledge of the relation between the effectors and their target molecules.

### Analyses of clinical datasets

mRNA expression of selected genes was analysed in the publicly available METABRIC^35^ and TCGA^36^ breast cancer cohorts using R (v4.5). Survival analyses were performed in the publicly available METABRIC cohort^35^ using overall survival (OS) and relapse-free survival (RFS). A signature score was calculated as the mean normalized *OSMR* and *ITGA5* co-expression, and patients were stratified into high and low groups using an optimal cut-off point based on the log-rank statistic.

Survival differences were assessed using Kaplan-Meier analysis and log-rank tests, and hazard ratios (HR) were estimated with Cox proportional hazards models. Analyses were conducted in all patients and in the basal-like subtype defined by PAM50 annotation^37^, using R (v4.5) and *survival* (v3.8.3), *survminer* (v0.5.1), *dplyr* (v1.1.4) and *ggplot2* (v4.0.1) R packages. Spearman rank gene correlations were computed between *OSM* or *OSMR* expression and all other genes in TCGA and METABRIC cohorts, including all samples. *P* values were adjusted for multiple testing using the Benjamini–Hochberg FDR method. Extracellular matrix–related genes were annotated using MatrisomeDB^38^. Correlation analyses were performed using cor.test() function from the *stats* R (v4.5) package.

### Statistics

Statistical analyses were performed using GraphPad Prism software and R (v4.5). For analyses involving two quantitative variables from patient datasets, Spearman’s correlation test was used. Associations between two qualitative variables were analysed using Fisher’s exact test. For comparisons between one quantitative and one qualitative variable, two-group comparisons were performed using a ratio paired *t*-test for paired *in vitro* experiments or a Welch-corrected *t*-test for *in vivo* experiments. Comparisons among more than two groups were performed using one-way ANOVA followed by Dunnett’s multiple-comparisons test or two-way ANOVA followed by Šidák’s multiple-comparisons test, as appropriate. Tumour growth curves were analysed using two-way ANOVA. Survival and tumour onset analyses were performed using Kaplan–Meier curves and compared using the log-rank (Mantel–Cox) test. For omics analyses, multiple tests were performed using Benjamini-Hochberg FDR method or multiple *t*-tests, differential features were visualized using volcano plots displaying –log_10_(FDR) or –log_10_(*P* value) versus log_2_ fold change. Pathway enrichment analyses were performed using Ingenuity Pathway Analysis and are reported as –log_10_(*P* value), z-score, and ratio. GSEA results are reported as normalized enrichment score (NES), gene ratio, and – log_10_(FDR). FDR or *P* values<0.05 were considered statistically significant. Unless otherwise stated, results are expressed as mean ± standard error of the mean (SEM).

### Study approval

All patient sample analyses have been performed in accordance with the Declaration of Helsinki. All procedures involving animals were performed with the approval of the Biogipuzkoa Animal Experimentation Committee and the Gipuzkoa Regional Government, according to European official regulations.

## Results

### OSM promotes cancer cell migration and invasion, and tumour growth *in vivo*

First, we studied the effects of Oncostatin M (OSM) cytokine signalling in breast cancer cells *in vitro* and *in vivo*. By combining exogenous treatment with recombinant human OSM and experiments with breast cancer cells ectopically overexpressing human OSM, we showed that OSM promoted migration and invasion of MDA-MB-231 cancer cells (**Figure 1A** and **Supplemental Figure 1A**), processes required for tumour growth and metastasis. We then generated tumours *in ovo* by injecting MDA-MB-231 cells in the chicken chorioallantoic membrane (CAM) model^21^ (**Supplemental Figure 1B**). In this model, OSM promoted tumour growth and increased distal cancer cell dissemination, detected by quantification of human *Alu* sequences in the distal CAM (**Supplemental Figure 1C-E**). In line with these results, orthotopic injection in nude mice of OSM-overexpressing cancer cells, alone or co-injected with cancer-associated fibroblasts (CAFs) (**Figure 1B-F**), led to accelerated tumour onset (**Supplemental Figure 2A**), increased tumour growth (**Figure 1C** and **F**) and micrometastasis, detected by the presence of human *Alu* sequences in lungs (**Figure 1D**). OSM-overexpressing tumours showed OSMR upregulation, supporting that the OSM pathway was active and functional^13^ (**Supplemental Figure 2B-C**). Importantly, therapeutic neutralization of OSM signalling with a receptor fusion protein (iOSM) in the immunocompetent 4T1 breast cancer model (**Figure 1G-H** and **Supplemental Figure 2D**) inhibited OSM effects on tumour growth (**Figure 1H**). Dose-dependent reduction of STAT3 activation by iOSM in OSM-treated 4T1 cells confirmed the efficacy of iOSM in blocking OSM downstream signalling (**Supplemental Figure 2E**).

**Figure 1:**
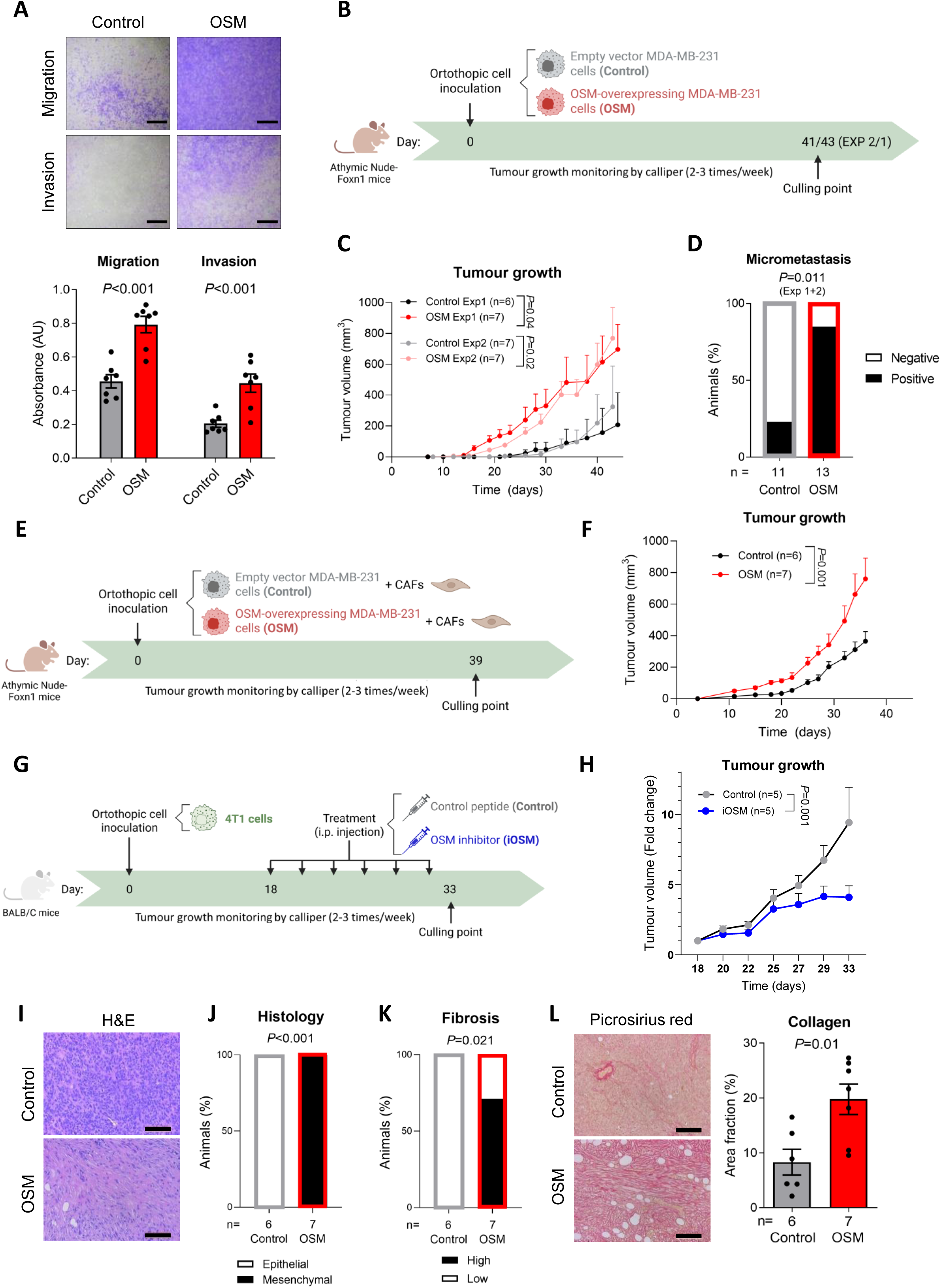
OSM promotes cancer cell migration, invasion, tumour growth and collagen remodelling. (**A**) Representative pictures (top) and quantification (bottom) of migration and invasion assays performed with control and OSM-overexpressing MDA-MB-231 cells. Scale bar: 1000 µm. (**B-F**) Schematic representations (**B, E**), tumour growth (**C, F**) and percentage of animals with lung micrometastases (**D**) from orthotopic xenograft experiments performed with control and OSM-overexpressing MDA-MB-231 cells injected alone (**B-D**) or with CAF-173 cells (**E-F**). (**G-H)** Schematic representation (**G**) and relative tumour growth since the start of the treatment (**H**) of orthotopic syngeneic 4T1 tumours treated with OSM inhibitor (iOSM) or control peptide. (**I**) Representative H&E pictures of control and OSM-overexpressing MDA-MB-231 orthotopic xenografts shown in C. Scale bar: 100 µm. (**J-K**) Pathological assessment of tumour histology (**J**), and fibrosis (**K**) from H&E images shown in **I**. (**L**) Picrosirius red representative stainings (left) and quantification (right) of control and OSM-overexpressing MDA-MB-231 orthotopic xenografts shown in **E**. Scale bar: 100 µm. Data represent mean ± SEM and *P* values were obtained using ratio-paired *t*-tests in **A**, two-way ANOVA in **C, F** and **H**, Fisher’s exact test in **D, J** and **K,** and *t*-test with Welch’s correction in **L**. In **A**, a total of 7 independent experiments were performed. AU: arbitrary units, Exp: experiment, H&E: haematoxylin and eosin.

### OSM activates ECM deposition and remodelling

We performed an in-depth characterization of control and OSM-activated orthotopic tumours to investigate the molecular mechanisms by which OSM promoted tumour progression. H&E histological analysis (**Figure 1I**) revealed that OSM-overexpressing tumours presented a mesenchymal-like phenotype (**Figure 1J**) and increased fibrosis (**Figure 1K**) compared to the control samples. This result was further validated by picrosirius red staining, which showed increased collagen deposition in OSM-activated tumours (**Figure 1L**). Interestingly, further analyses revealed that OSM signalling activation also increased proline hydroxylation in cancer cells (**Supplemental Figure 2F**), a process that directly impacts on collagen structure and promotes breast cancer progression^39^.

Proteomic and transcriptomic analyses of control and OSM-activated tumours revealed regulation of multiple ECM and matrisome-associated genes and proteins by OSM (colour-coded in the indicated figures), including collagens, glycoproteins, proteoglycans, and ECM regulators such as LOX, PLOD2, TGM2, serpins, cathepsins and collagen prolyl hydroxylases 3 and 4 (**Figure 2A-B** and **Supplemental Tables 5-6**).

**Figure 2:**
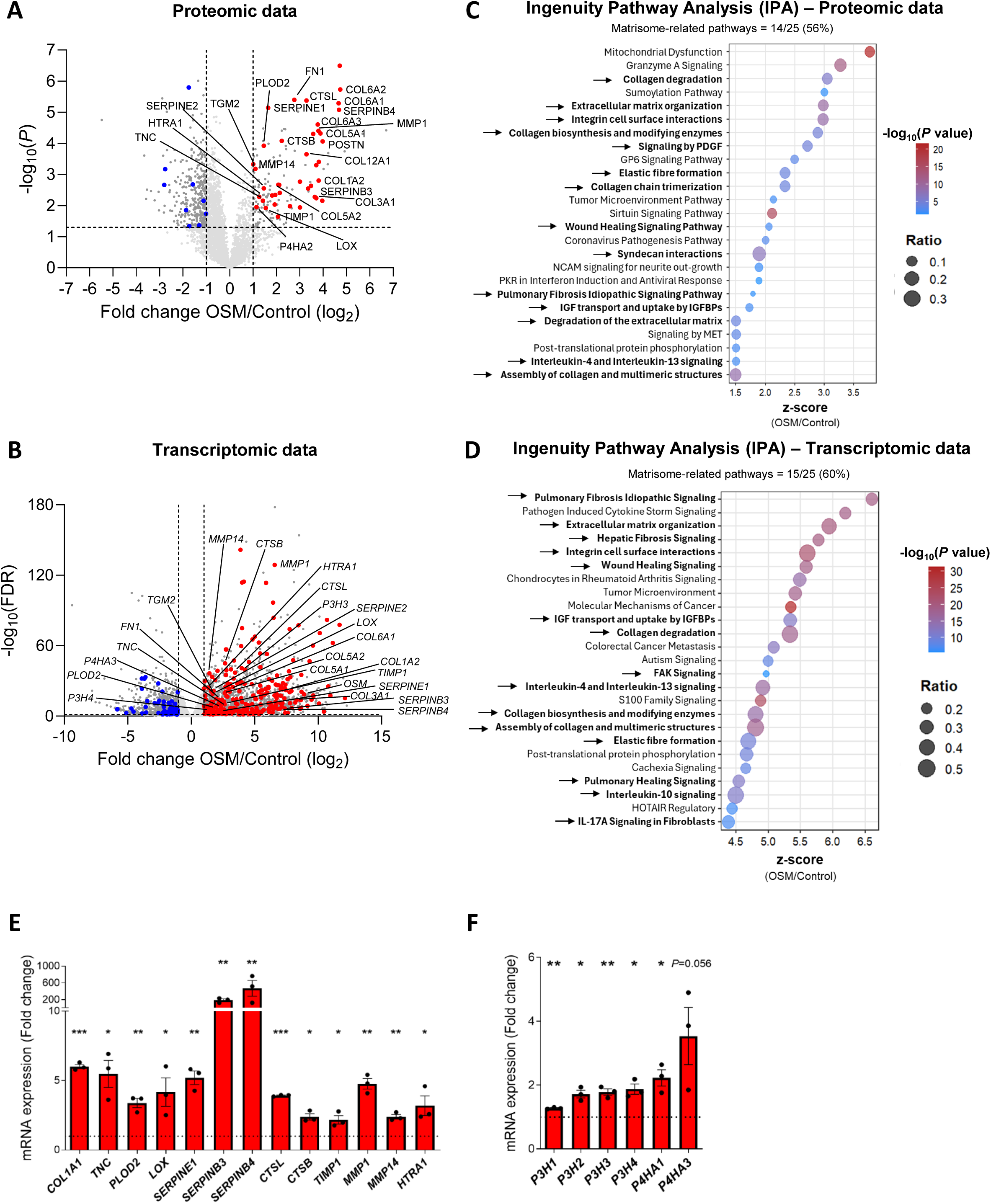
OSM promotes the expression of multiple ECM molecules and ECM regulators. (**A-B**) Volcano plots of differentially expressed proteins (**A**) and genes (**B**) in proteomic (**A**) and transcriptomic (**B**) analyses of orthotopic tumours derived from control or OSM-overexpressing MDA-MB-231 cells injected with CAF-173 cells. Selected cut-offs (*P* value or FDR<0.05 and Fold change >|2|) are shown in dashed lines. Colours show matrisome-related significant genes/proteins in red (upregulated) or blue (downregulated), while genes/proteins not related to the matrisome are shown in dark grey when significantly deregulated and light grey when non-significant. (**C-D**) Top pathways identified by Ingenuity Pathway analyses (IPA) from deregulated proteins in **C** or genes in **D**. Matrisome-related pathways are shown in bold and indicated with arrows. Selected cut-offs were *P* value<0.05 and z-score >|1.5|. The ratio shows the coverage of detected genes/proteins in each pathway. (**E-F**) mRNA expression levels of the indicated ECM-related (**E**) or prolyl hydroxylases (**F**) analysed by RT-qPCR in OSM-overexpressing cells grown in 3D spheres *in vitro* compared to its corresponding controls. Data represent mean ± SEM. FDR or *P* values were obtained using multiple testing by the Benjamini–Hochberg method or *t*-tests respectively in **A**, **B**, **C** and **D**, and using ratio-paired *t*-tests in **E** and **F**. AU: Arbitrary units. FDR: False Discovery Rate. ECM: Extracellular matrix. (*) *P*<0.05, (**) *P*<0.01, (***) *P*<0.001.

Importantly, 26% of proteins (40 out of 153, **Figure 2A** and **Supplemental Figure 3A**) and 12% of genes (292 out of 2432, **Figure 2B**) upregulated by OSM are considered core matrisome or matrisome-associated proteins^38^. Ingenuity Pathway Analysis (IPA) of the proteomic data of OSM-activated tumours confirmed that, among the top 25 pathways enriched by OSM, 14 (56%) were associated with the matrisome, such as ECM remodelling, integrin interactions, wound healing and collagen remodelling, biosynthesis and degradation (**Figure 2C** and **Supplemental Table 7**). This phenotype was confirmed by IPA analysis of RNA-seq data (**Figure 2D** and **Supplemental Table 8**), as well as by pathway analysis of transcriptomic data derived from OSM-activated cancer cells *in vitro* and tumours inoculated without CAFs (**Supplemental Figure 3B-E** and **Supplemental Tables 9-10**). We further confirmed that OSM increased gene expression of key ECM components and regulators, such as *COL1A1*, *TNC, PLOD2, LOX*, serpins, cathepsins, metalloproteinases and proteases, in 3D heterospheroids formed by MDA-MB-231 cancer cells and CAFs (**Figure 2E**). In line with our data supporting that OSM increased prolyl hydroxylation, OSM also increased the expression of different members of the prolyl hydroxylases 3 and 4 families (**Figure 2F**), key proteins in collagen prolyl hydroxylation^40^. Taken together, these data indicate that OSM:OSMR signalling promotes ECM deposition and remodelling, and the expression of ECM regulators.

### ITGA5 is an important mediator of OSM pro-malignant effects

Integrins and integrin-mediated adhesions have long been recognized as the main molecular link between cells and the ECM, and are known to connect signals between the extracellular environment and intracellular pathways in a bidirectional way^1,3^. We analysed the modulation of integrins by OSM in our transcriptomic and proteomic data and observed that many integrins were upregulated by OSM in cancer cells and in tumour samples (**Figure 3A-C**). To check the relevance of these findings in the clinical setting, we analysed the correlation between *OSM* and *OSMR* genes with all the α (ITGA) and β (ITGB) integrin subunits in breast cancer tumour samples from patients included in the TCGA and METABRIC cohorts (**Figure 3D** and **Supplemental Tables 11-12**). The expression of *OSM* and *OSMR* positively correlated with the expression of most integrins in TCGA and METABRIC datasets.

**Figure 3:**
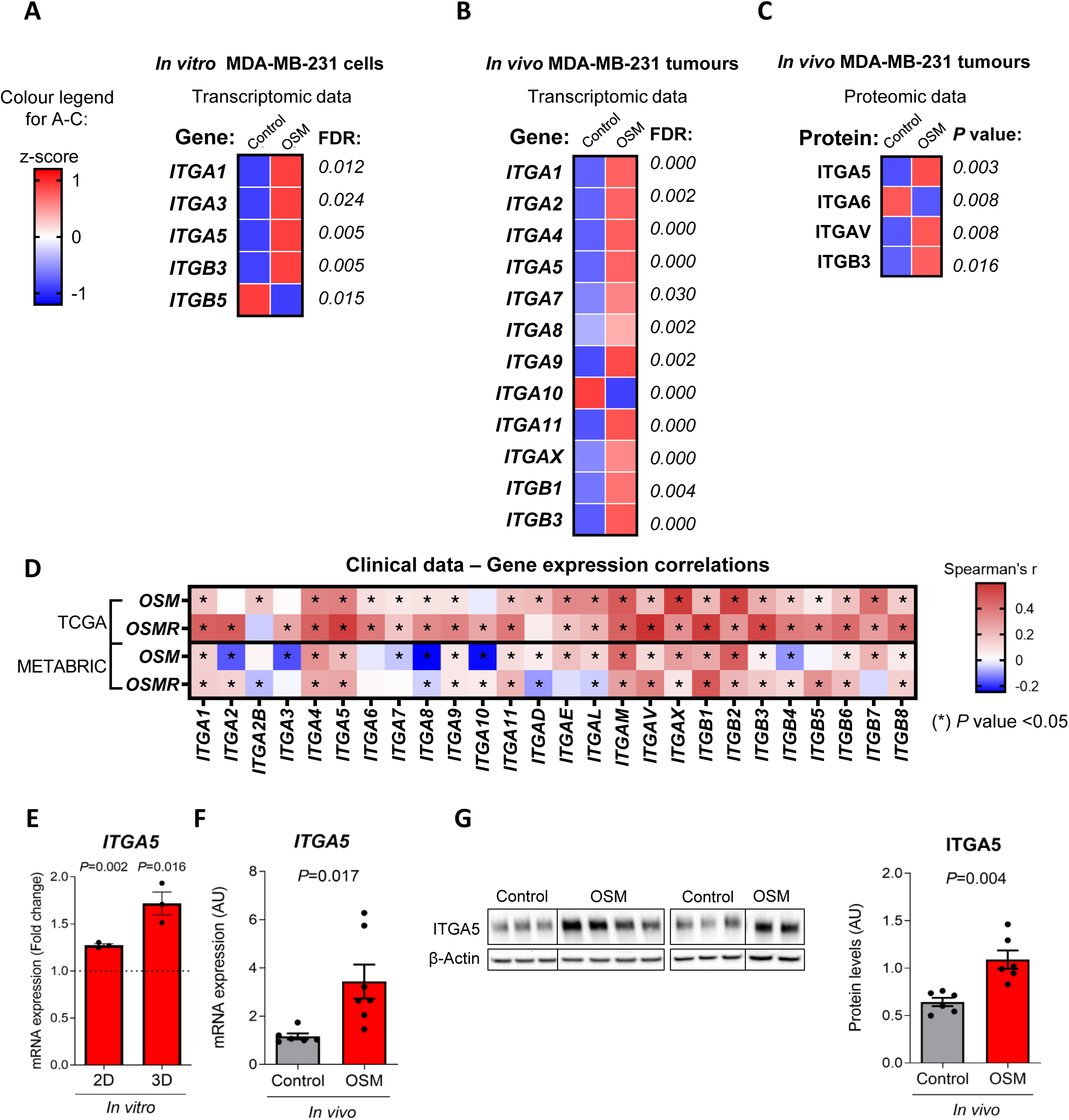
OSM promotes the expression of integrins. (**A-C**) Heatmaps showing normalized mRNA (**A-B**) and protein (**C**) expression of integrins induced by OSM in transcriptomic (**A-B**) and proteomic (**C**) data from control and OSM-overexpressing cells *in vitro* (**A**) and *in vivo* in orthotopic tumours derived from control or OSM-overexpressing MDA-MB-231 cells injected with CAF-173 cells (**B-C**). (**D**) Correlation of *OSM* and *OSMR* mRNA levels with α and β integrin subunits in breast cancer samples from TCGA (n = 1111) and METABRIC (n = 1980) cohorts. (**E-G**) mRNA (**E-F**) and protein (**G**) levels of ITGA5 analysed by RT-qPCR (**E-F**) and Western Blot (**G**) in OSM- overexpressing cells, compared to their corresponding controls, grown in 2D and 3D spheres *in vitro* (**E**) or in orthotopic tumours derived from control or OSM-overexpressing MDA-MB-231 cells injected with CAF-173 cells (**F-G**). Representative Western Blot panels are shown in **G**. FDR and *P* values were obtained using multiple testing by the Benjamini–Hochberg method or *t*-tests respectively in **A-C**, ratio-paired *t*-tests in **E**, and *t*-test with Welch’s correction in **F** and **G**. Spearman’s correlation analysis was performed in **D** with associated *P* values calculated with Spearman’s rank correlation coefficients. Data represent mean ± SEM. 4 independent experiments were performed in **A**, and 3 in **E**. 4 animals per group were analysed in **B-C,** and 6 or 7 in **F-G**. FDR: False Discovery Rate. (*) *P*<0.05.

Among all these integrins, we focused on ITGA5, as it is the integrin whose expression was most strongly correlated with OSM and OSMR in breast cancer clinical samples and which was consistently induced by OSM in all our transcriptomic and proteomic datasets (**Figure 3A-D**). First, we validated the transcriptomic and proteomic data confirming OSM-mediated ITGA5 upregulation in breast cancer cells *in vitro* (2D and 3D heterospheroids) and tumours by RT-qPCR and Western Blot analyses, respectively (**Figures 3E-G**). Finally, to further confirm that OSM-mediated ITGA5 upregulation was occurring via OSMR, we overexpressed *OSMR* in SK-BR-3 cells, a breast cancer cell line that does not express *OSMR* endogenously. We observed that, in comparison with control cells, *OSMR*-overexpressing SK-BR-3 cells showed a clear increase of *ITGA5* expression upon treatment with OSM (**Supplemental Figures 4A-B**).

To study if the increase of *ITGA5* expression by OSM in cancer cells was functionally relevant, we silenced it with small interfering RNA (siRNA) and subjected MDA-MB-231 cells to migration and invasion assays (**Figure 4A**). *ITGA5* was efficiently silenced in both control and OSM overexpressing cells (**Supplemental Figure 4C-D**). While ITGA5 silencing did not affect control cells, it impaired the increased migration and invasion induced by OSM (**Figure 4A**). In addition, *ITGA5* silencing also resulted in smaller tumours in the CAM model (**Supplemental Figure 4E**).

**Figure 4:**
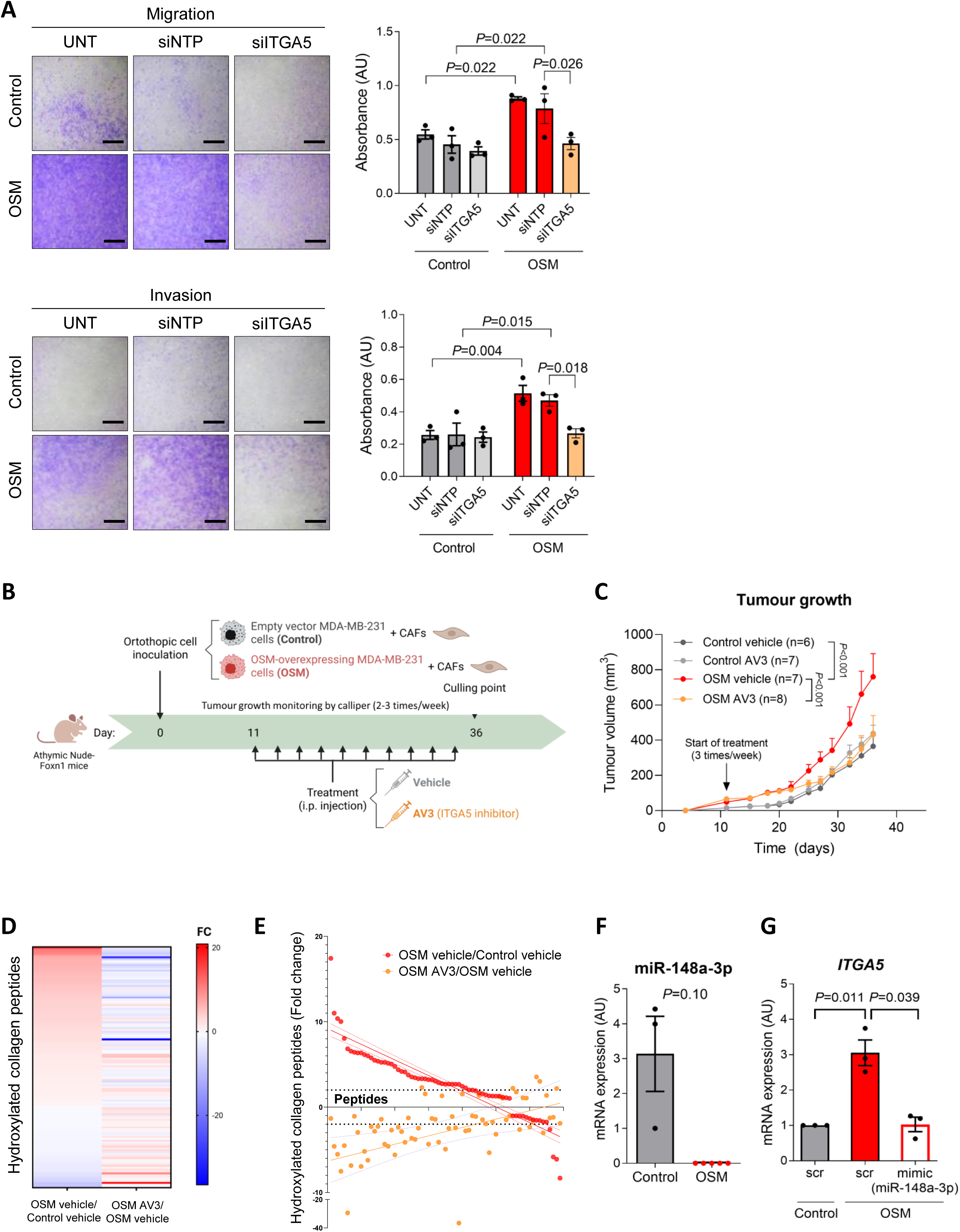
ITGA5 is an important mediator of OSM-pro-malignant effects in cancer cells. (**A**) Migration (top panels) and invasion (bottom panels) assays performed with control and OSM-overexpressing MDA-MB-231 cells transfected with siRNAs targeting ITGA5 (siITGA5), non-targeting control siRNAs (non-targeting pool, siNTP) or untransfected (UNT). Representative pictures of crystal-violet-stained transwells are shown (left panels), and the corresponding quantification of migration and invasion assays is shown in the right panels. Scale bar: 1000 µm. (**B-C**) Schematic representation (**B**) and tumour growth (**C**) of orthotopic xenograft experiments performed with control and OSM-overexpressing MDA-MB-231 cells co-injected with CAF-173 cells and treated with ITGA5 inhibitor (AV3) or vehicle. The tumour growth of control vehicle and OSM vehicle tumours is also shown in Figure 1F. (**D-E**) Heatmap (**D**) and relative quantification (**E**) of hydroxylated collagen peptides in tumours from panel B. (**F**) miR-148a-3p expression in control and OSM-activated tumours from experiment 2 in Figure 1B. (**G**) *ITGA5* expression in control and OSM-overexpressing MDA-MB-231 cells upon transfection with the mimic of miR-148a-3p or scramble (scr) miRNA. Data represent mean ± SEM and *P* values were obtained using Šidák’s multiple comparisons test (**A**), two-way ANOVA (**C**), *t*-test with Welch’s correction (**F**) and ratio-paired *t*-tests in **G**. 3 independent experiments were performed in **A** and in **G**. 4 tumours per group were analysed in **D**, **E** and 3-5 in **F**.

We then wondered if therapeutic ITGA5 inhibition could halt the pro-tumour effects of OSM *in vivo*. Control or OSM-stimulated cells were orthotopically co-injected with CAFs, and tumours were treated with the AV3 peptide ITGA5 inhibitor^20^ (**Figure 4B**).

The effect of OSM in tumour growth was completely abrogated when ITGA5 was blocked with AV3 (**Figure 4C**). At the molecular level, ITGA5 inhibition reduced OSM-induced collagen hydroxylation, as analysed in proteomic data of these tumours (**Figure 4D-E** and **Supplemental Table 13**). In agreement with the data shown in **Supplemental Figure 2F**, OSM signalling increased proline hydroxylation in 62% of the detected collagen peptides (FC ≥2 in 43 out of 69). Importantly, ITGA5 inhibition reversed the effect of OSM as it reduced the hydroxylation of 72% of these peptides (FC ≥-2, 31 out of 43 peptides increased by OSM) (**Figure 4D-E** and **Supplemental Table 13**).

In order to dissect the molecular mechanisms by which OSM promoted ITGA5 expression and activation, we interrogated the microRNA Data Integration Portal (mirDIP v.5.0.2.3)^41^ to find miRNAs able to target *ITGA5*, and only miRNAs with a very high score (top 1%) were considered. The miRNA with the highest score was hsa-miR-148a-3p, which, interestingly, had been identified as a STAT3 transcriptional target in ChiP-seq experiments (ENCODE^42^) and in the Eukaryotic Promoter Database^43^. Our analyses showed that OSM downregulated the expression of miR-148a-3p in MDA-MB-231 cancer cells *in vitro* (**Supplemental Figure 4F**) and *in vivo* (**Figure 4F**), with no detectable expression in OSM-activated tumours. To determine if miR-148a-3p downregulation was involved in ITGA5 induction by OSM, breast cancer cells were transfected with a synthetic miR-148a-3p (mimic) or a negative control miRNA (scr) (**Supplemental Figure 4G**). ITGA5 induction by OSM was blocked when cells were transfected with the miRNA mimic (**Figure 4G**). In summary, our data showed that OSM signalling promotes integrin expression in breast cancer cells, through miR-148- 3p downregulation, and that ITGA5 mediates OSM’s pro-malignant effects in our models.

### OSMR-ITGA5 axis is increased in tumours from basal breast cancer patients where it associates with decreased survival

We then analysed breast cancer clinical samples to confirm the relevance of our findings in the clinical setting. Gene expression data from the publicly available METABRIC database showed that the expression of *OSM*, *OSMR* and *ITGA5,* the main components of the pro-malignant axis identified herein, was upregulated in breast cancer patients from the basal subtype (**Figure 5A**). In line with our *in vitro* and *in vivo* results, GSEA of the genes that positively correlated with *OSMR* in clinical tumour samples, demonstrated that those genes were associated to gene signatures related to matrisome, ECM regulation and integrin signalling in samples from basal breast cancer patients, as well as in samples from all breast cancer subtypes (**Figure 5B**, **Supplemental Figure 5A** and **Supplemental Tables 4, 14** and **15**).

**Figure 5:**
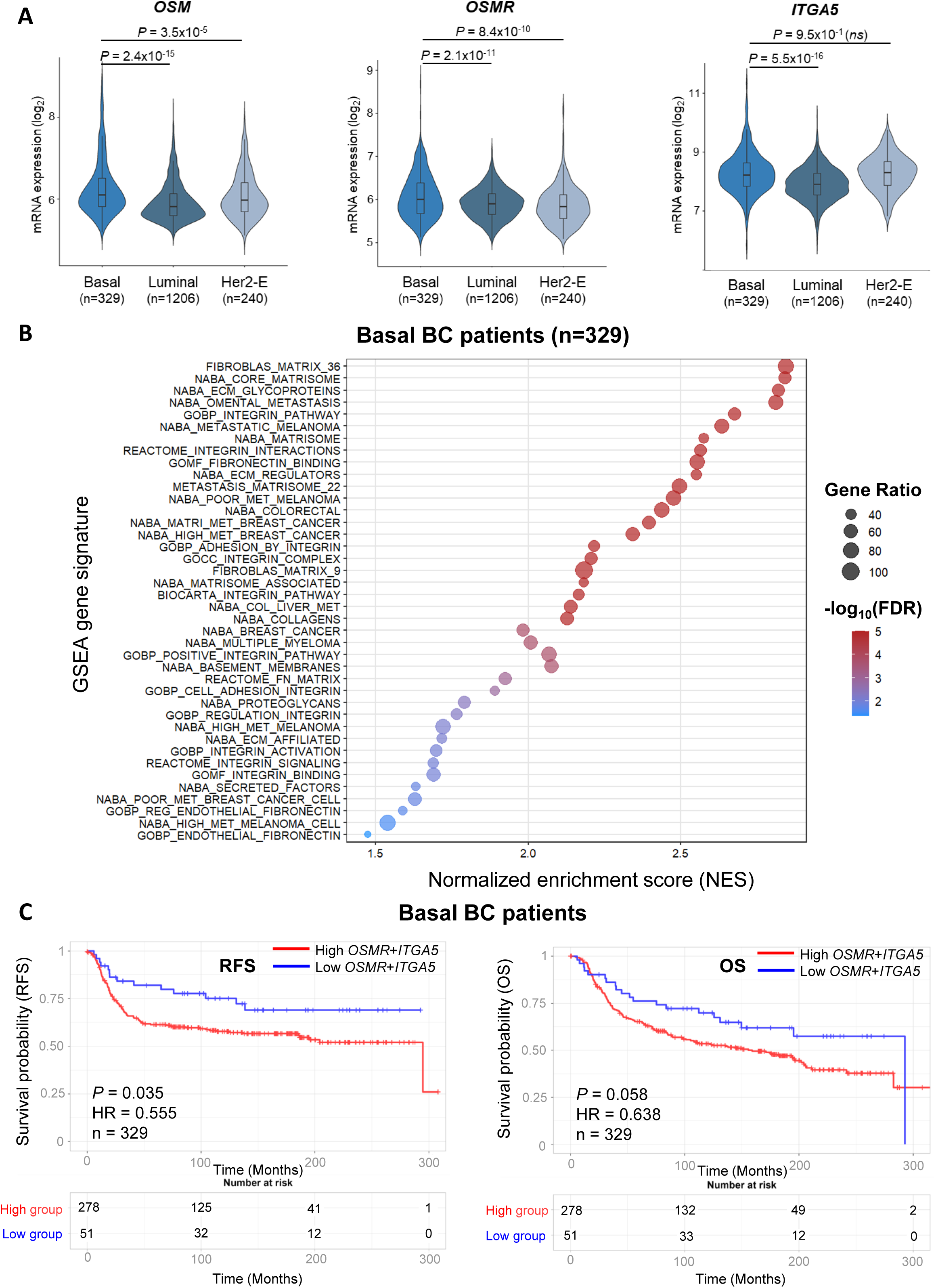
*OSM-OSMR-ITGA5* is increased in tumours from basal breast cancer patients and associates with reduced survival. (**A**) *OSM*, *OSMR* and *ITGA5* mRNA expression in breast cancer clinical samples from patients included in the METABRIC dataset, stratified by breast cancer subtypes. (**B**) Gene Set Enrichment Analysis (GSEA) of ECM and integrin-related gene signatures in transcriptomic data of basal tumours from the METABRIC cohort. Genes were ordered according to their Spearman’s correlation coefficient with the *OSMR* gene. Significance was considered with FDR values below 0.05 and Normalized Enrichment Score (NES) above 1.5. Gene Ratio represents the percentage of coverage of detected genes in each signature. (**C**) Kaplan-Meier curves showing recurrence-free survival (RFS) and overall survival (OS) for breast cancer patients of the basal subtype included in the METABRIC cohort, according to the tumour mRNA expression of *OSMR* and *ITGA5*. *P* values were determined using the Mantel-Cox test. Hazard ratios (HR) were estimated using Cox proportional hazards models. In **A**, *P* values were obtained using one-way ANOVA Dunnett’s post hoc tests, considering the basal subtype group as reference. In **B**, false discovery rate (FDR) q- values were calculated using permutation-based testing to estimate the proportion of false positives among gene sets with similar or greater enrichment scores. BC: breast cancer. ECM: extracellular matrix. Her2-E: HER2 enriched. ns: non-significant.

We then analysed if *OSMR* and *ITGA5* co-expression could have an impact in patient survival. Interestingly, high *OSMR-ITGA5* co-expression was associated with reduced recurrence-free (RFS) and overall survival (OS), both when analysing only basal breast cancer patients (**Figure 5C**) or patients from all breast cancer subtypes (**Supplemental Figure 5B**) included in the METABRIC cohort. Importantly, lower hazard ratios (HRs) were obtained when analyses were performed in the basal breast cancer cohort supporting the relevance of this axis in this breast cancer subtype. Taken together, our data support that OSMR-ITGA5 signalling axis associates with gene signatures related to ECM remodelling and with decreased survival in breast cancer patients, and that this pathway is particularly relevant in patients from the basal subtype.

## Discussion

It is nowadays well accepted that the ECM plays a critical role in cancer progression, as it has been linked to all stages of cancer from tumour initiation to metastasis^1,44,45^.

However, ECM-targeting therapies have so far shown limited success in clinical trials, despite promising results in preclinical *in vivo* studies^44^. Understanding the dynamic reciprocal nature of cancer cell–ECM interactions and the upstream regulators of ECM remodelling is key to design effective therapeutic strategies to block tumour progression. Breast cancer (BC) is a suitable model for addressing these research questions as it is considered a desmoplastic tumour, characterized by an excessive or abnormal deposition of ECM, a dense fibrotic stroma and fibroblast expansion and activation^1^.

Here we show that the cytokine OSM is a master regulator of ECM remodelling in breast cancer by promoting ECM deposition, activating the expression of ECM regulators and cross-linkers, and of integrins. The ability to remodel the ECM has been classically attributed to cancer-associated fibroblasts (CAFs). We and others recently demonstrated that OSM activates CAFs by promoting fibroblast activation protein alpha (FAP) expression, CAF proliferation, and chemokine secretion in breast and pancreatic cancers^13,15^. OSM also promotes the expression of ECM components and remodelling factors in chondrocytes and lung, dermal and cardiac fibroblasts, contributing to fibrosis, joint inflammation and atherosclerosis^11,46–50^.

Importantly, ECM can also be produced by cancer cells, and increased ECM production by tumour cells promotes metastasis and correlates with poor patient survival^5,51^. In line with this, our results reveal a key role for cancer cells as producers of ECM genes and regulators, although the contribution of CAFs cannot be excluded in some of our models. Our data show that OSM promotes collagen deposition and increases the expression of tenascin and fibronectin at protein and mRNA level. It also promotes collagen hydroxylation, and the expression of many ECM remodelling proteins and cross-linking enzymes such as lysyl oxydase (LOX), prolyl hydroxylases, serpins, cathepsins and PLOD2. Aberrant collagen deposition and crosslinking in the breast are associated with increased mammographic density and a higher risk of breast cancer^52^. Cross-linking enzymes, along with collagen’s hydroxylation state and fibronectin cross-linking by transglutaminase 2 (TGM2) have been associated to tumour progression and metastasis in many tumour types^53^. We and others previously showed that OSM promotes the expression of fibronectin (FN1) and TGM2 in breast and cervical cancer cells *in vitro*^54,55^. In summary, our data shows that OSM cytokines activates ECM deposition and remodelling and promotes the expression of many ECM regulators.

Among the main ECM receptors (integrins, discoidin domain receptors and syndecans^44^), we observed a general upregulation of integrins, which expression consistently correlated with *OSM* and *OSMR* in BC clinical samples (**Figure 3**). Many integrins, such as ITGAV, ITGA4, ITGA5,ITGA6, ITGB1 or ITGB3 are overexpressed in various cancer types and activate tumour cell invasion and metastasis upon binding to the ECM^1^. They are actually involved in every step of tumour progression, from primary tumour growth to metastatic dissemination^2,3^. Our data show that integrin α5 (ITGA5) is an important mediator of OSM pro-malignant effects (**Figure 4**). ITGA5, together with ITGB1, forms the α5β1 heterodimer, which binds specifically to fibronectin^56^. Both α5β1 integrin and fibronectin have been associated with metastasis in breast cancer^57,58^. Fibronectin has been shown to regulate the composition and stability of the ECM and is required for the deposition and organization of fibrillar collagen I^59^, suggesting that the effects of OSM in the different ECM proteins may be connected. In addition, TGM2, the co-receptor for fibronectin together with α5β1^60^, is a matrix-crosslinking enzyme of different ECM proteins such as fibronectin itself, but also fibrinogens, heparan sulphate proteoglycans and collagen VI^61^. In line with this complex crosstalk between the different ECM proteins and regulators, our data show that ITGA5 blocking reverts the effects of OSM on collagen hydroxylation (**Figure 4 C-D**). Other reports have pointed to the effects of ITGA5 inhibition in collagen. For example, α5-blocking antibodies and peptides and the ITGA5 inhibitor dioscin, reduced collagen synthesis in blood vessels, pancreatic cancer models and hepatic stellate cells^20,62,63^. Therefore, our data supports that ITGA5 mediates, at least in part, the effects of OSM in tumour growth, acting as a sensor of the ECM and participating in collagen remodelling.

As ECM-targeting therapies have been of great interest but yet remained unsuccessful in clinical trials^44^, alternative approaches would be to target how cells sense this pathological matrix, by inhibiting integrins. However, clinical trials testing the efficacy of integrin-targeting drugs in solid tumours have shown limited success so far. Poor pharmacokinetics of some of these drugs, lack of biomarkers to assess efficacy, and the dynamic activation and wide heterogeneity of integrins are some of the factors that could explain integrin therapy failure in the past^44^. For instance, Volociximab is an anti-α5β1 integrin antibody that showed promising results in Phase 1b trials^64^, but was not effective in monotherapy in a Phase II clinical trial including chemotherapy-resistant ovarian and peritoneal cancer patients^65^. Dual or multi-targeting therapies may be a more rational choice to target integrins in cancer. New ways to block specific integrins are currently being pursued. As an example, AV3, the peptidomimetic ITGA5 inhibitor tested herein, potentiated the efficacy of chemotherapy in pancreatic cancer models^20^ and integrin α5β1 targeting antibodies increased the efficacy of immunotherapy in breast cancer models^62^.

The results presented herein support that an alternative approach to therapeutically target the malignant ECM in breast cancer, would be to target factors, such as OSM, that promote the tumour pathological matrix. Antibodies targeting OSM signalling are in clinical development for the treatment of inflammatory diseases. In particular, anti-OSM (GSK2330811) and anti-OSMR (vixarelimab) humanized antibodies are being tested in Phase 2 clinical trials for the treatment of systemic sclerosis, prurigo nodularis or ulcerative colitis (NCT06693908, NCT05785624)^66,67^. The knowledge generated on these inflammatory diseases in which antibodies targeting cytokines and associated receptors are currently being used in the clinic^6,8^, could help the translation of OSM-blocking antibodies to the cancer arena. In fact, we previously showed that the GSK2330811 anti-OSM antibody inhibits invasion, angiogenesis and metastasis in cervical cancer^16^ and we show herein that OSM inhibition with a receptor fusion protein reduced tumour growth in an immunocompetent mouse model of triple negative breast cancer.

In conclusion, our results prove that the cytokine OSM activates ECM deposition and remodelling, resulting in a tumour-associated aberrant matrix. Alterations in ECM are mainly sensed by integrins and our data reveal that ITGA5 inhibition, by both genetic and pharmacological means, abrogates the pro-malignant effects of OSM in breast cancer models (**Figure 6**). We also showed that *OSM* and *OSMR* correlated with matrisome genes and integrins in human breast cancer samples and that co-expression of OSMR and ITGA5 associated with decreased survival. In line with this, *OSM*, *OSMR* and *ITGA5* genes have been included in gene signatures associated with bad prognosis in glioblastoma and head and neck squamous cell carcinoma^68,69^. Considering the therapeutic efficacy of OSM and ITGA5 inhibitors in mouse breast cancer models presented in this manuscript, our research sheds light on the potential use of this axis as a therapeutic target in human breast cancer in the future.

**Figure 6:**
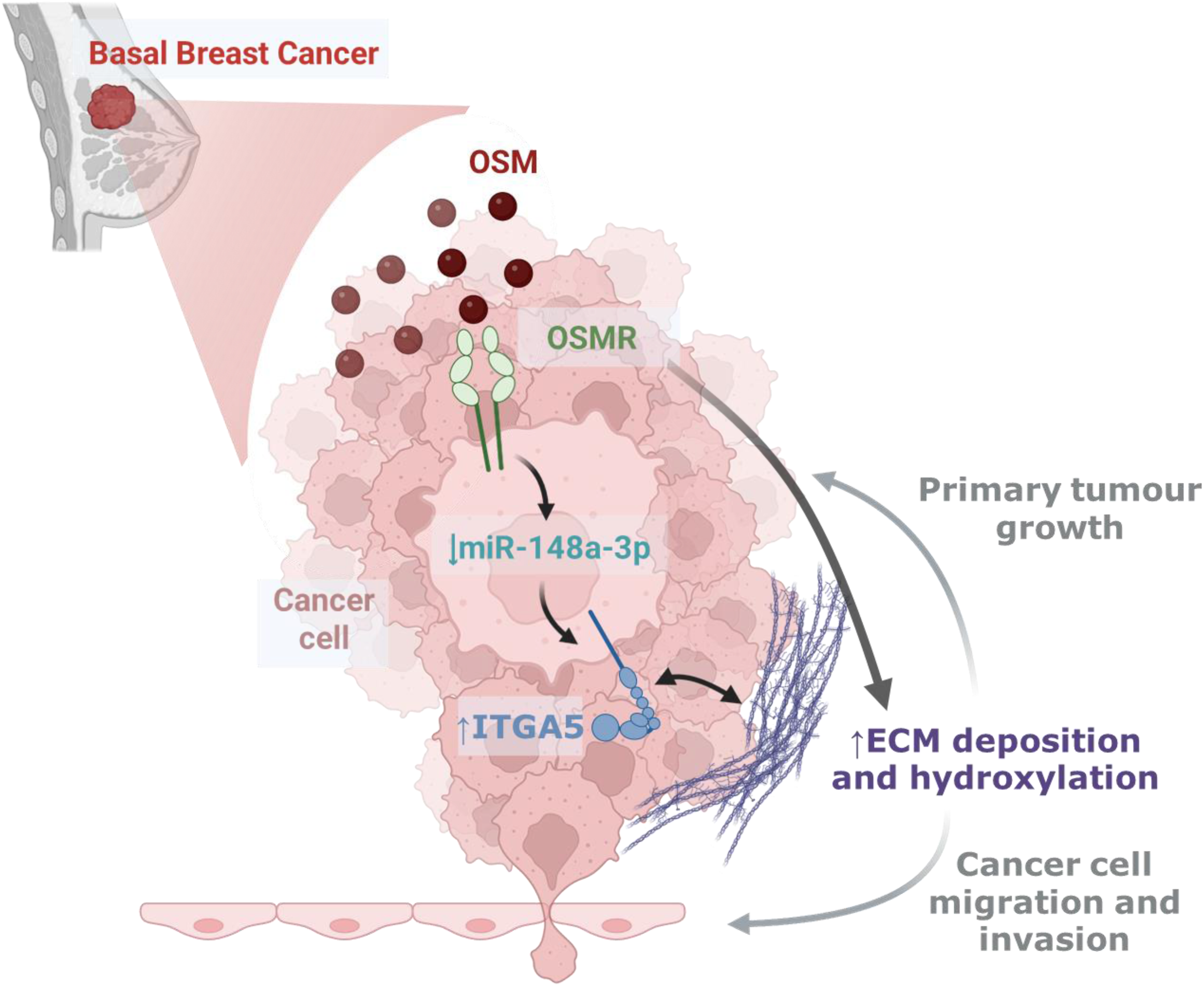
Graphical Abstract. Oncostatin M cytokine promotes breast cancer progression by remodelling the extracellular matrix and activating integrin signalling in cancer cells. OSM: Oncostatin M. OSMR: Oncostatin M Receptor. ITGA5: integrin alpha-5. ECM: extracellular matrix. miR: microRNA.

## Abbreviations

BC,: Breast cancer
CAF,: Cancer-associated fibroblast
CAM,: Chorioallantoic membrane
ECM,: Extracellular matrix
iOSM,: Oncostatin M inhibitor
LC-MS/MS: Liquid chromatography–tandem mass spectrometry
OSM,: Oncostatin M
OSMR,: Oncostatin M receptor
TME,: Tumour microenvironment
TNBC,: Triple-negative breast cancer

## Acknowledgements

We are grateful to the members of our laboratories for critical discussion of this work and to the Genomics and Histology Platforms and Animal Facility of the Biogipuzkoa Health Research Institute for technical assistance and advice. We thank Dr Paloma Bragado (UCM, Madrid, Spain), Dr Patricia Fernandez-Nogueira (University of Barcelona, Barcelona, Canada) for providing the human cancer-associated fibroblasts and Prof Sarah-Maria Fendt for her critical insights on experimental design.

## Funding

This work was funded by Spanish Ministry of Science and Innovation - ISCIII (PI18/00458, PI21/01208 and FI19/00193), co-funded by the European Union; Spanish Ministry of Science and Innovation (CNS2023-145020, CPP2022-009535 and PID2024-158438OB-I00); BBVA Leonardo Award (LEO23-2-10814-BBM-TRA-229); Fundación Asociación Española Contra el Cáncer (LABAE247490MUNO); Fundación FERO; Fundación Gangoiti; Fundación SEOM (Beca SEOM-FESEO 2021); Basque Department of Industry, Tourism and Trade (Elkartek); Basque Department of Health (20200111040), and Ikerbasque Basque Research Foundation. The group also received funds from the breast cancer patient’s charity Katxalin and from Roche Farma S.A. AA, AMAraujo, AA and UAH were funded by Basque Government Doctoral Training Grants. SM is funded by a Sara Borrell Fellowship from ISCIII (CD23/00059), co-funded by the European Union. JILV was funded by an AECC PhD Fellowship. SS is funded by the Special Research Fund starting grant of the KU Leuven (STG/22/034). MMC is funded by Ikerbasque.

## Author Contributions

AA, PA, SM and AMAraujo performed all the cellular and molecular experiments. SM performed the CAM model. AA, PA, SM, AMAraujo, JILV, ZT and MMC performed the animal experiments. JMF performed histology experiments and JMF and MR analysed mouse histopathology. AA, PA, UAH and PD performed bioinformatic analyses with help from DO. PA and SS performed collagen hydroxylation experiments. AMAransay performed and analysed RNA-seq experiments. MA and FE performed and analysed proteomic experiments. GMN provided the OSM inhibitor. JP provided the ITGA5 inhibitor, contributed with experimental design and helped with supervision of the project. JP and MMC provided funds. AA and MMC designed the study. MMC supervised and led the project. AA, PA, SM, UAH, PD and MMC analysed the data and wrote the manuscript. AMAraujo, JILV, GMN, AMAransay, MA, SS and JP edited the manuscript, and all authors gave final approval to the submitted version of the manuscript.

## Competing interests

Authors declare that they have no competing interests.

## Data and materials availability

The mRNA and protein datasets generated during the current study by microarray) and proteomics (MSV000101746) are available in the GEO and MassIVE repositories. GSE195787 https://www.ncbi.nlm.nih.gov/geo/query/acc.cgi?acc=GSE195787 GSE331508 https://www.ncbi.nlm.nih.gov/geo/query/acc.cgi GSE328180 https://www.ncbi.nlm.nih.gov/geo/query/acc.cgi MSV000101746 https://massive.ucsd.edu/ProteoSAFe/dataset.jsp?task=71608b1d9f764416a732b8c9ec3d6003

Source data for uncropped Western blots are provided in **Supplemental Figure 6**. All other data files supporting the findings of this study are available from the corresponding author upon reasonable request.

## Notes

### Competing Interest Statement

The authors have declared no competing interest.

