## Supplementary Figures and supp figure and table legends for "Oncostatin M cytokine promotes breast cancer progression by remodelling the extracellular matrix and activating integrin signalling in cancer cells"

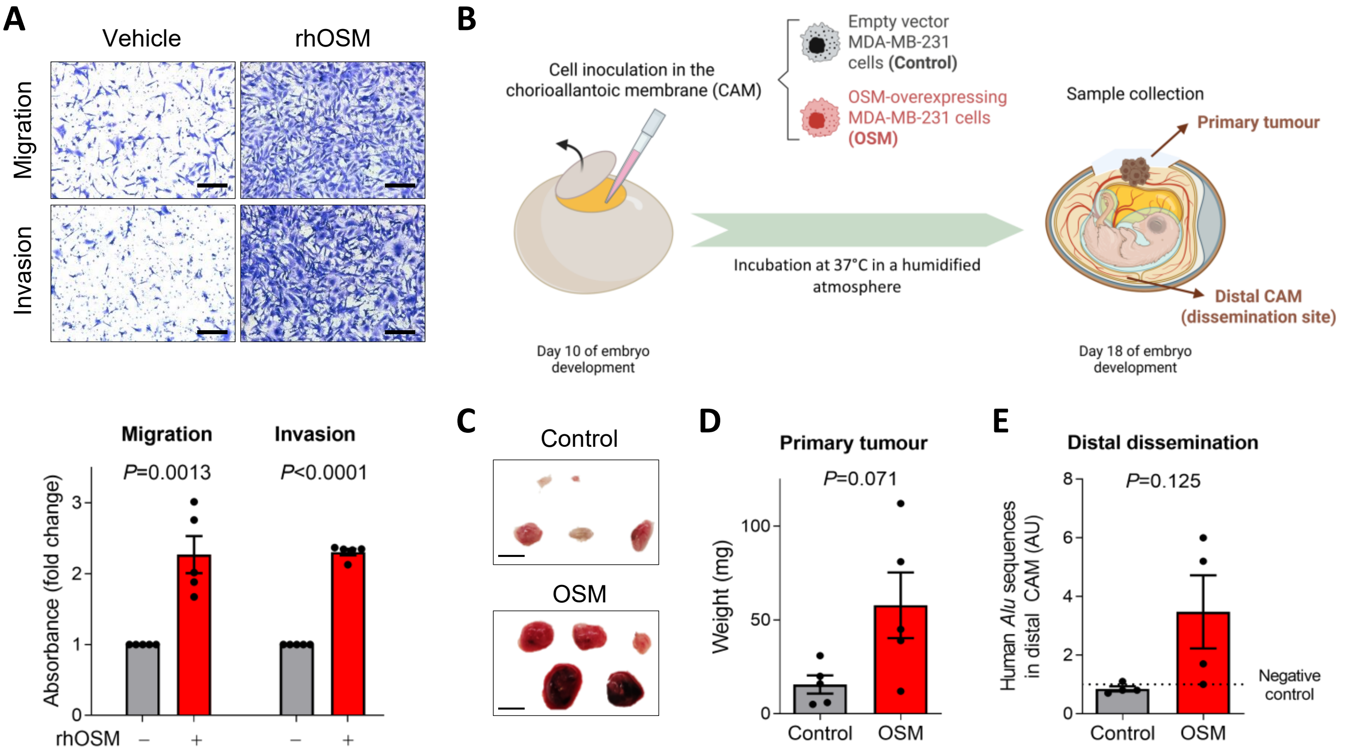
A
Vehicle
rhOSM
Migration
Invasion
Migration
Invasion
P=0.0013
P<0.0001
Absorbance (fold change)
rhOSM
B
Cell inoculation in the chorioallantoic membrane (CAM)
Empty vector MDA-MB-231 cells (Control)
OSM-overexpressing MDA-MB-231 cells (OSM)
Sample collection
Primary tumour
Distal CAM (dissemination site)
Incubation at 37°C in a humidified atmosphere
Day 10 of embryo development
Day 18 of embryo development
C
Control
OSM
D
Primary tumour
P=0.071
Weight (mg)
Control
OSM
E
Distal dissemination
P=0.125
Human Alu sequences in distal CAM (AU)
Negative control
Control
OSM

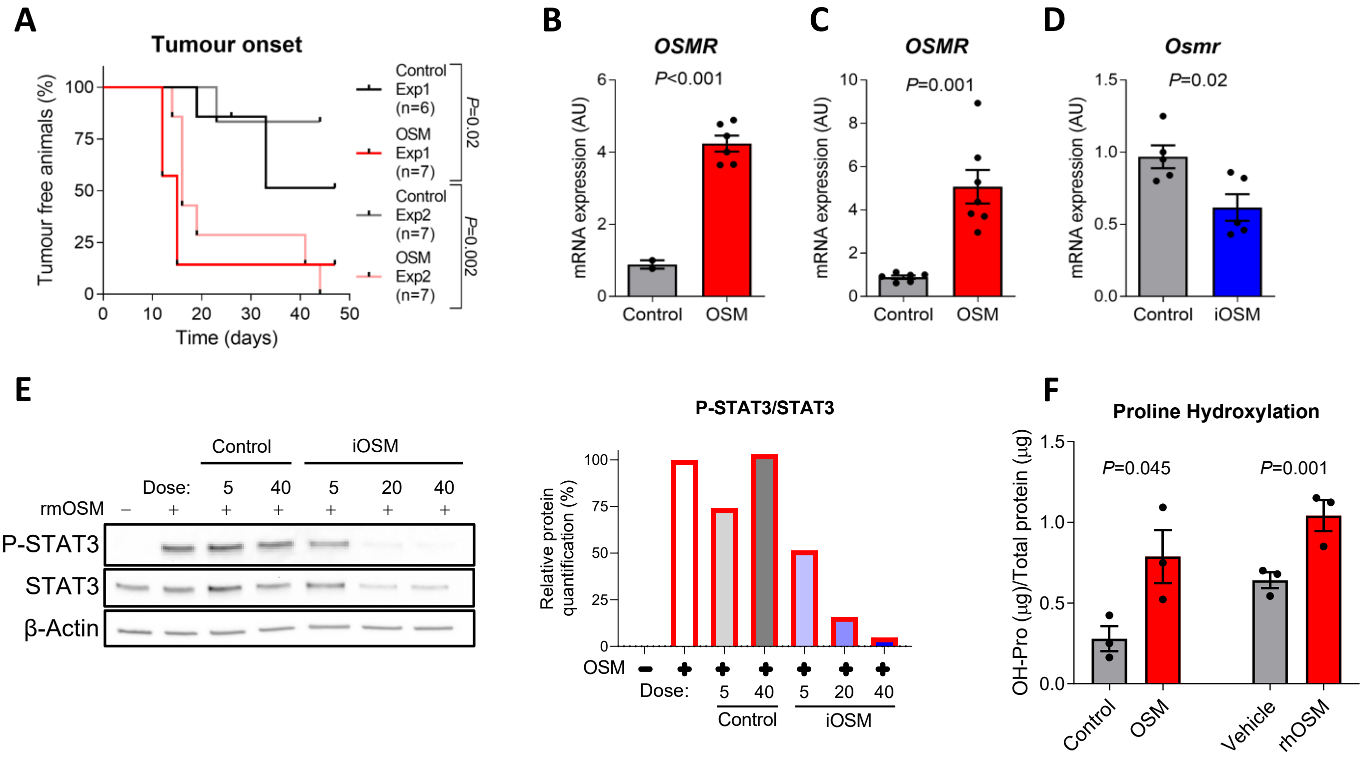
A
Tumour onset
Tumour free animals (%)
Time (days)
Control Exp1 (n=6)
OSM Exp1 (n=7)
Control Exp2 (n=7)
OSM Exp2 (n=7)
P=0.02
P=0.002
B
OSMR
P<0.001
mRNA expression (AU)
Control
OSM
C
OSMR
P=0.001
mRNA expression (AU)
Control
OSM
D
Osmr
P=0.02
mRNA expression (AU)
Control
iOSM
E
Control
iOSM
Dose: 5 40 5 20 40
rmOSM – + + + + + +
P-STAT3
STAT3
β-Actin
P-STAT3/STAT3
Relative protein quantification (%)
OSM
Dose: 5 40 5 20 40
Control
iOSM
F
Proline Hydroxylation
OH-Pro (µg)/Total protein (µg)
P=0.045
P=0.001
Control
OSM
Vehicle
rhOSM

**A**

| Matrisome division | Matrisome category | Enriched in OSM-activated tumours (n) | Enriched in OSM-activated tumours (%) |
|---|---|---|---|
| **Core matrisome** | | | |
| | Collagens | 8 | 5.2 |
| | ECM Glycoproteins | 12 | 7.8 |
| | Proteoglycans | 3 | 2.0 |
| **Matrisome-associated** | | | |
| | ECM regulators | 14 | 9.2 |
| | ECM-affiliated proteins | 1 | 0.7 |
| | Secreted factors | 2 | 1.3 |
| | **Total** | 40/153 | 26.2 |

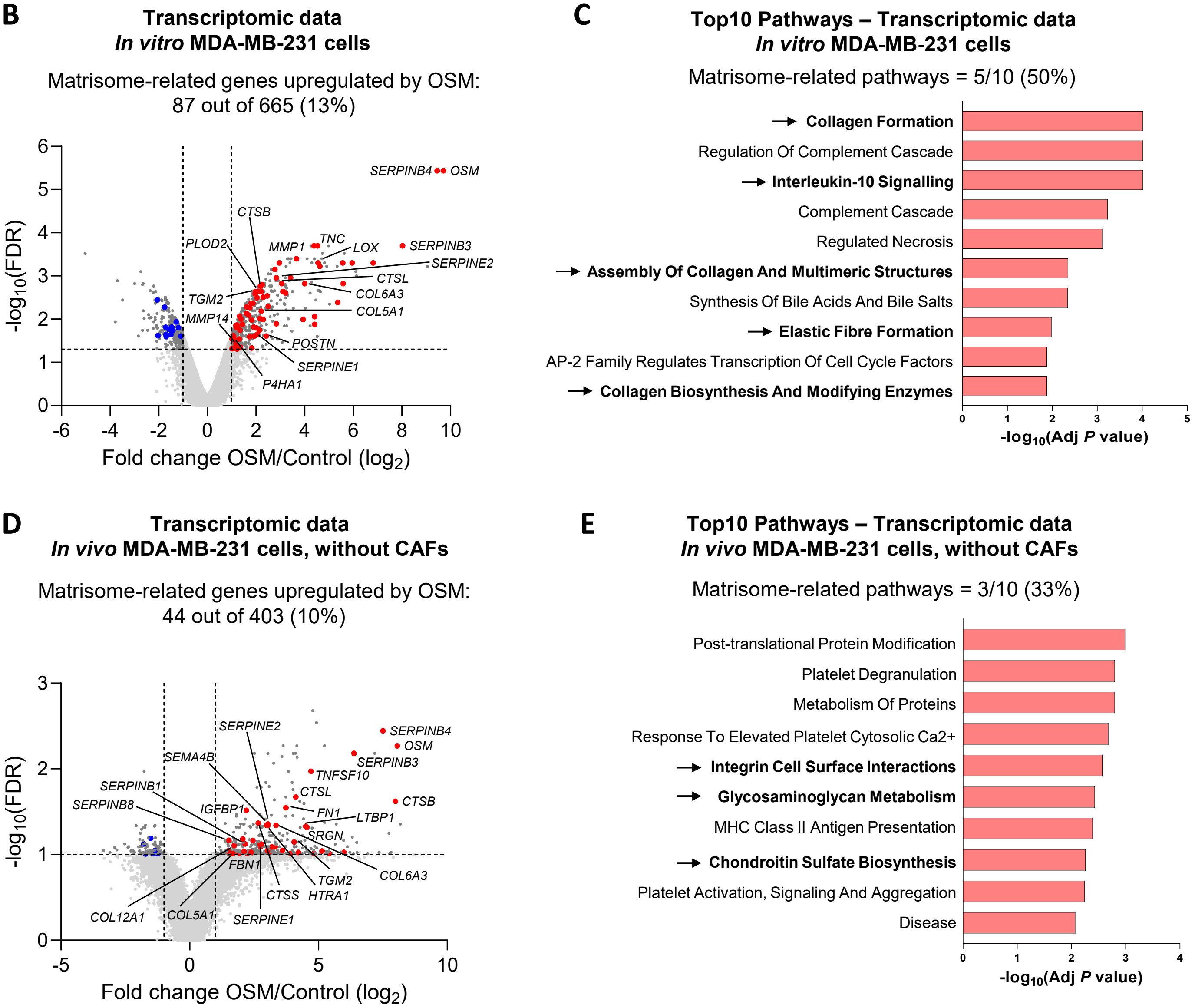

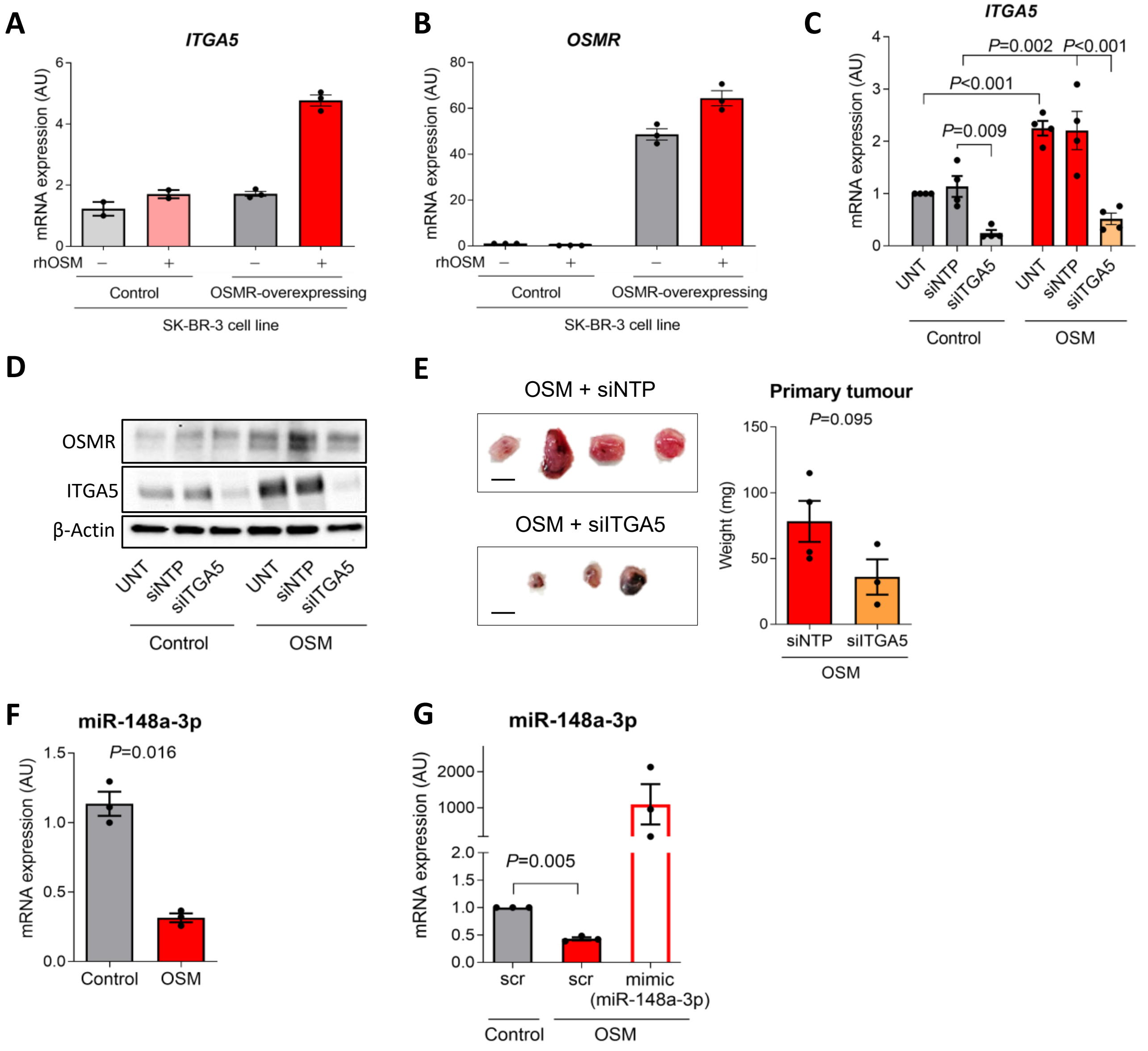

A
ITGA5
mRNA expression (AU)
rhOSM
Control
OSMR-overexpressing
SK-BR-3 cell line
B
OSMR
mRNA expression (AU)
rhOSM
Control
OSMR-overexpressing
SK-BR-3 cell line
C
ITGA5
mRNA expression (AU)
P=0.002
P<0.001
P<0.001
P=0.009
UNT
siNTP
siITGA5
UNT
siNTP
siITGA5
Control
OSM
D
OSMR
ITGA5
β-Actin
UNT
siNTP
siITGA5
UNT
siNTP
siITGA5
Control
OSM
E
OSM + siNTP
OSM + siITGA5
Primary tumour
P=0.095
Weight (mg)
siNTP
siITGA5
OSM
F
miR-148a-3p
P=0.016
mRNA expression (AU)
Control
OSM
G
miR-148a-3p
mRNA expression (AU)
P=0.005
scr
scr
mimic
(miR-148a-3p)
Control
OSM

**A** **All BC patients (n=1980)**

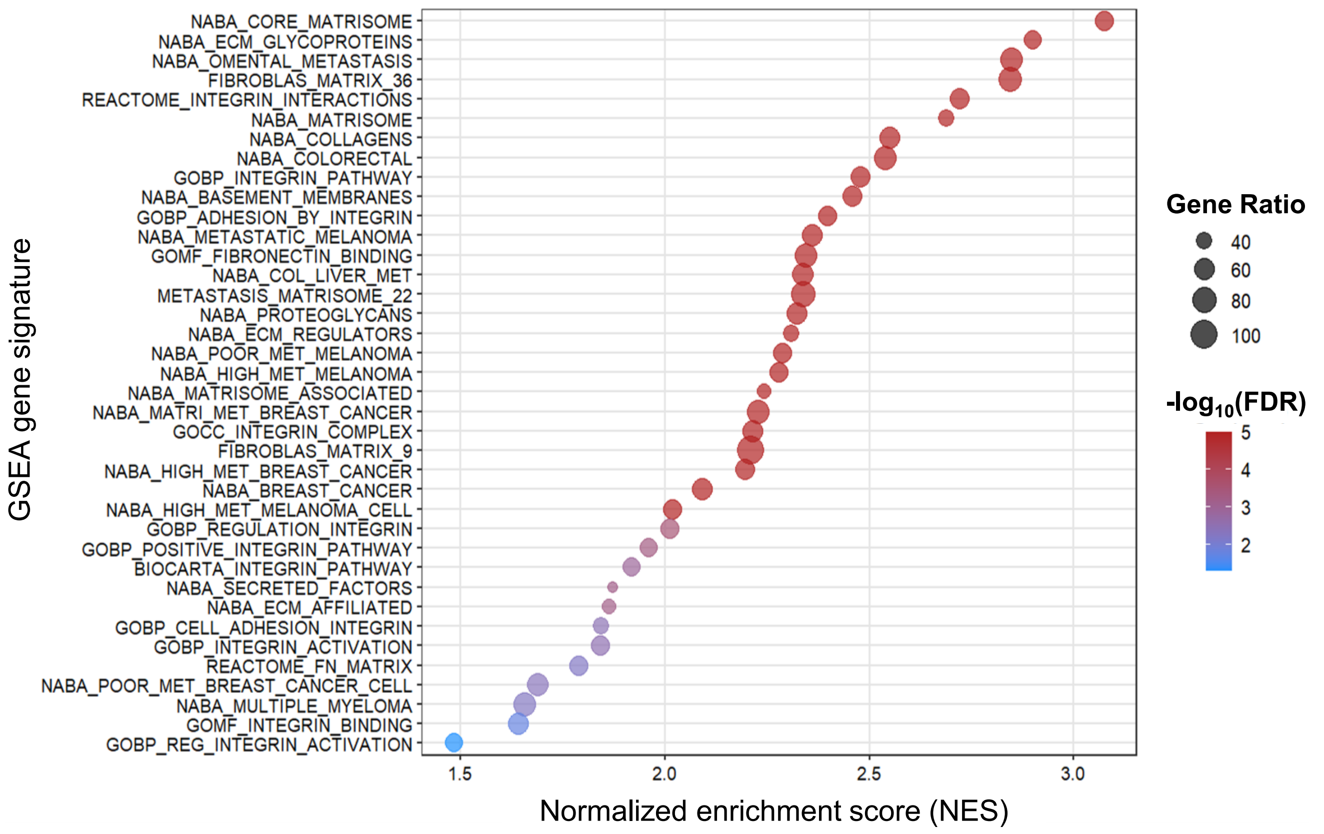


**B** **All BC patients**

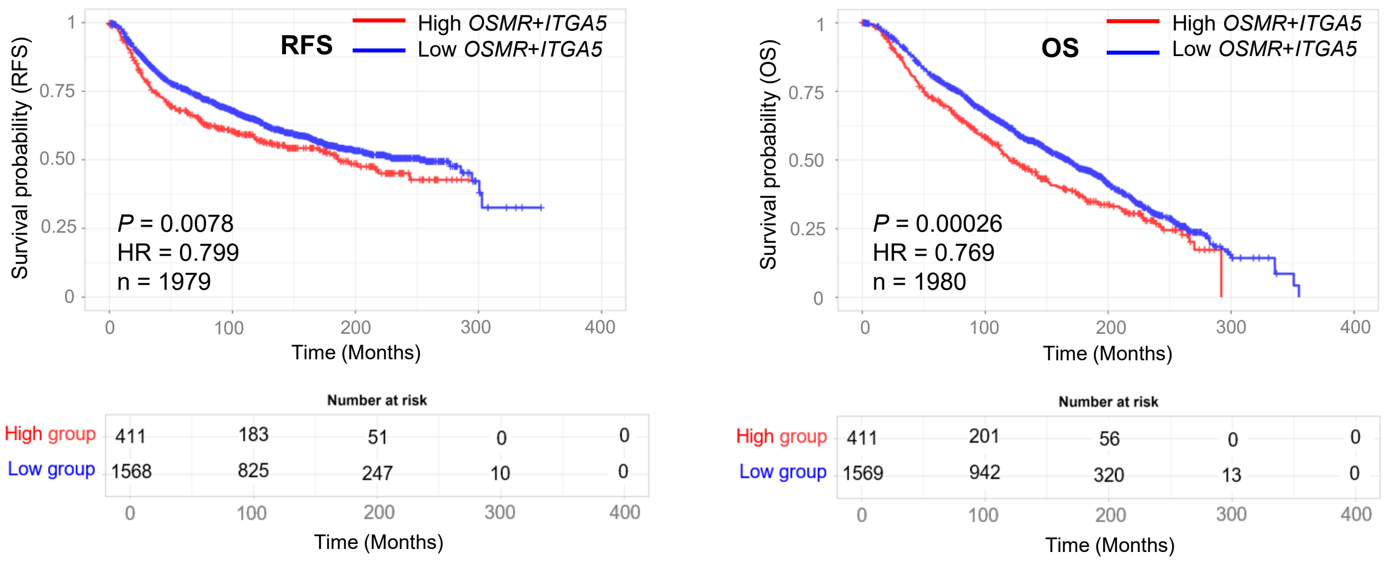

**A** Full unedited gels for Figure 3, panel G

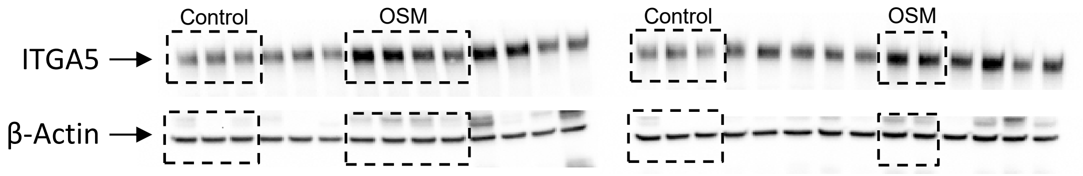


**B** Full unedited gels for Supplemental Figure 2, panel E

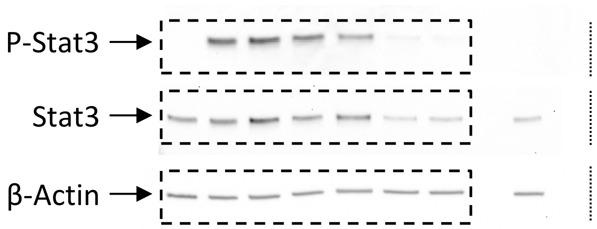


**C** Full unedited gels for Supplemental Figure 4, panel D

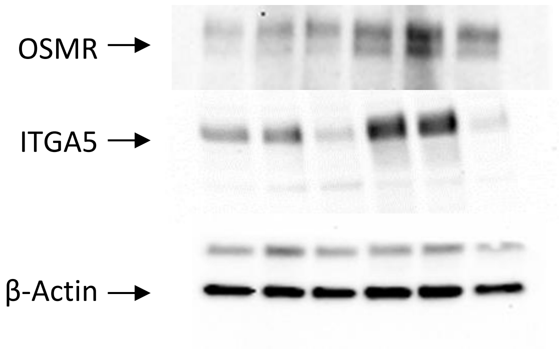

**Supplemental Figure Legends**

**Figure S1: Effects of OSM *in vitro* and in the *in ovo* CAM model.** (**A**) Representative pictures (top) and quantification (bottom) of migration and invasion assays performed with MDA-MB-231 cells treated with vehicle or recombinant human OSM (rhOSM). Scale bar: 200 µm (**B-E**) Schematic representation (**B**), representative pictures (**C**), tumour growth (**D**) and dissemination (**E**) of the *in ovo* CAM tumour model performed with control and OSM-overexpressing MDA-MB-231 cells. Scale bar: 0.5 cm. Data represent mean ± SEM, and *P* values were obtained using ratio-paired *t*-tests in **A** and *t*-test with Welch's correction in **D** and **E**. A total of 5 independent experiments were performed in **A**, and 5 eggs per experimental condition were inoculated in **B-E**, 4 per condition being analysed for distal dissemination (**E**). AU: Arbitrary units.

**Figure S2: Effects of OSM and OSM inhibitor in preclinical breast cancer *in vivo* and *in vitro* models.** (**A-B**) Tumour onset (**A**) and OSMR mRNA levels in tumours (**B**) of orthotopic xenograft experiments performed with control and OSM-overexpressing MDA-MB-231 cells injected alone. (**C**) OSMR mRNA levels in tumours of orthotopic xenograft experiments performed with control and OSM-overexpressing MDA-MB-231 cells injected with CAF-173 cells. (**D**) OSMR mRNA levels in orthotopic syngeneic 4T1 tumours treated with OSM inhibitor (iOSM). (**E**) Representative Western Blots (left) and densitometric quantification (right) of P-STAT3, STAT3 and β-Actin protein levels in 4T1 cells treated with 5 ng/ml recombinant murine OSM (rmOSM) and increasing doses of OSM inhibitor (iOSM), expressed as iOSM: OSM molar ratios. (**F**) Quantification of hydroxylated prolines (OH-Pro) in collagens measured in control and

OSM-overexpressing MDA-MB-231 cells (left) and in MDA-MB-231 cells treated with recombinant human OSM (rhOSM) or vehicle (right). Data represent mean ± SEM and *P* values were obtained using log-rank (Mantel-Cox) test in **A**, *t*-test with Welch's correction in **B**, **C** and **D** and ratio-paired *t*-test in **F**. Data in panel **E** shows a representative blot and quantification from one independent experiment. Exp: Experiment. AU: Arbitrary units.

**Figure S3: Proteomic and transcriptomic data of OSM-activated MDA-MB-231 cells *in vitro* and *in vivo*.** (**A**) Proportions of ECM-related proteins in OSM-activated orthotopic tumours derived from control or OSM-overexpressing MDA-MB-231 cells injected with CAF-173 cells. (**B-E**) Volcano plots of differentially expressed genes (**B, D**) and top 10 enriched pathways (**C, E**) in transcriptomic analyses of control and OSM-overexpressing MDA-MB-231 cells *in vitro* (**B, C**) and *in vivo* (**D, E**). In **B, D**, colours show matrisome-related significant genes in red (upregulated) or blue (downregulated), while genes not related to the matrisome are shown in dark grey when significantly deregulated and light grey when non-significant. In **C, D**, matrisome-related pathways are shown in bold and indicated with arrows. Selected cut-offs in Volcano Plots (FDR<0.05 in **B** and FDR<0.1 in **D**, and Fold change >|2|) are shown in dashed lines. *P* values or FDRs were obtained using multiple testing by the Benjamini–Hochberg method or *t*-tests respectively. Adj: Adjusted. CAFs: Cancer associated fibroblasts. ECM: Extracellular matrix.

**Figure S4: Role of ITGA5 and miR-148a-3p as mediators of OSM-effects.** (**A-B**) *ITGA5* (**A**) and *OSMR* (**B**) mRNA expression levels analysed by RT-qPCR in control and OSMR-overexpressing SK-BR-3 cells treated with vehicle or rhOSM. Graphs represent technical replicates from one representative experiment. (**C-D**) ITGA5 mRNA (**C**) and protein (**D**) levels, analysed by RT-qPCR and Western Blot respectively, of control and OSM-overexpressing MDA-MB-231 cells transfected with siRNAs targeting ITGA5 (siITGA5), non-targeting control siRNAs (non-targeting pool, siNTP) or untransfected (UNT). (**E**) Representative pictures (left) and tumour growth (right) of the *in ovo* CAM tumour model performed with OSM-overexpressing MDA-MB-231 cells transfected with siRNAs targeting ITGA5 (siITGA5) or non-targeting control siRNAs (non-targeting pool, siNTP). Scale bar: 0.5 cm. (**F-G**) miR-148a-3p expression in control and OSM-overexpressing MDA-MB-231 cells in basal conditions (**F**), and upon transfection with the mimic of miR-148a-3p or scramble (scr) miRNA (**G**). Data represent mean ± SEM and *P* values were obtained using two-way ANOVA and Šidák's multiple comparisons test in **C**, *t*-test with Welch's correction in **E** and ratio-paired *t*-tests in **F** and **G**. A total of 4 independent experiments were performed in **C** and **D** and 3 experiments in **F** and **G**.  4 eggs per condition were inoculated in **E**, and 1 egg in the OSM + siITGA5 condition did not generate any tumour.

**Figure S5: Correlation of *OSMR* with genes related to the matrisome and association of *OSMR-ITGA5* axis with reduced survival in the METABRIC breast cancer cohort.** (**A**) Gene Set Enrichment Analysis (GSEA) of ECM and integrin-related gene signatures in transcriptomic data of all tumours from the METABRIC cohort. Genes were ordered according to their Spearman's correlation coefficient with the *OSMR* gene. Significance was considered with FDR values below 0.05 and Normalized Enrichment Score (NES) above 1.5. Gene Ratio represents the percentage of coverage of detected genes in each signature. False discovery rate (FDR) q-values were calculated using permutation-based testing to estimate the proportion of false positives among gene sets with similar or greater enrichment scores. (**B**) Kaplan-Meier curves showing recurrence-free (RFS) and overall survival (OS) for all breast cancer patients included in the METABRIC cohort, according to the tumour mRNA expression of *OSMR* and *ITGA5*. *P* value was determined using the Mantel-Cox test. Hazard ratios (HR) were estimated using Cox proportional hazards models. BC: breast cancer.

**Figure S6: Full unedited gels for Western blots shown in main and supplemental figures.**

**Supplemental table legends**

**Supplemental Table 1:** References for all reagents used in the Materials and Methods section.

**Supplemental Table 2:** Sequence of RT-qPCR primers, Taqman probes and synthetic siRNA and miRNAs used in this study. HK: housekeeping.

**Supplemental Table 3:** List of antibodies used for Western Blotting in this study.

**Supplemental Table 4:** ECM and integrin-related gene-signatures used in this study.

**Supplemental Table 5:** List of detected proteins in the proteomic analyses of control and OSM-activated tumours derived from orthotopic injection of control and OSM-overexpressing MDA-MB-231 cells co-injected with CAF-173 cells. The ratio of abundance between the OSM and control groups is shown for each protein. For proteins belonging to the matrisome, the matrisome division and category are shown. *P* values were obtained using multiple comparisons from 4 samples per experimental group.

**Supplemental Table 6:** List of detected genes in the RNA-seq analyses of control and OSM-activated tumours derived from orthotopic injection of control and OSM-overexpressing MDA-MB-231 cells co-injected with CAF-173 cells. The ratio of abundance and FDR between the OSM and control groups is shown for each gene. For genes belonging to the matrisome, the matrisome division and category are shown. FDR values were obtained using multiple comparisons from 4 samples per experimental group.

**Supplemental Table 7:** Ingenuity Pathway Analysis (IPA) of the proteomic data of control and OSM-activated tumours derived from orthotopic injection of control and OSM-overexpressing MDA-MB-231 cells co-injected with CAF-173 cells. Selected cut-offs were *P* value<0.05 and z-score >|1.5|.

**Supplemental Table 8:** Ingenuity Pathway Analysis (IPA) of the transcriptomic data of control and OSM-activated tumours derived from orthotopic injection of control and OSM-overexpressing MDA-MB-231 cells co-injected with CAF-173 cells. Selected cut-offs were *P* value<0.05 and z-score >|1.5|.

**Supplemental Table 9:** List of detected genes in the microarray analysis of control and OSM-overexpressing MDA-MB-231 cells *in vitro*. The ratio of abundance and FDR between the OSM and control groups is shown for each gene. For genes belonging to the matrisome, the matrisome division and category are shown. Fold change and FDR values were obtained using multiple comparisons from 4 samples per experimental group.

**Supplemental Table 10:** List of detected genes in the microarray analyses of control and OSM-activated tumours derived from orthotopic injection of control and OSM-overexpressing MDA-MB-231 cells. The ratio of abundance and FDR between the OSM and control groups is shown for each gene. For genes belonging to the matrisome, the matrisome division and category are shown. Fold change and FDR values were obtained using multiple comparisons from 4 samples per experimental group.

**Supplemental Table 11:** Correlation analysis of *OSM* and *OSMR* mRNA levels with all the α (ITGA) and β (ITGB) integrin subunits in breast cancer clinical samples from patients included in the TCGA cohort (n= 1111). Spearman’s correlation coefficients were obtained along with *P* values and q values.

**Supplemental Table 12:** Correlation analysis of OSM and OSMR mRNA levels with all the α (ITGA) and β (ITGB) integrin subunits in breast cancer clinical samples from patients included in the METABRIC cohort (n= 1980). Spearman’s correlation coefficients were obtained along with *P* values and q values.

**Supplemental Table 13:** List of hydroxylated collagen peptides from proteomic analyses of tumours derived from control and OSM-overexpressing MDA-MB-231 cells co-injected with CAF-173 cells and treated with ITGA5 inhibitor (AV3) or vehicle. The ratio of abundance among groups is shown for each peptide.

**Supplemental Table 14:** Correlation analysis of *OSMR* mRNA levels with all genes included in the transcriptomic data of breast cancer clinical samples from patients of the basal subtype included in the METABRIC cohort (n= 329). For genes belonging to the matrisome, the matrisome division and category are shown. Spearman’s correlation coefficients, *P* values and FDR values are shown.

**Supplemental Table 15:** Correlation analysis of *OSMR* mRNA levels with all genes included in the transcriptomic data of breast cancer clinical samples from all patients included in the METABRIC cohort (n= 1980). For genes belonging to the matrisome, the matrisome division and category are shown. Spearman’s correlation coefficients, *P* values and FDR values are shown.
